# Capillary electrophoresis mass spectrometry identifies new isomers of inositol pyrophosphates in mammalian tissues

**DOI:** 10.1101/2022.09.14.507917

**Authors:** Danye Qiu, Chunfang Gu, Guizhen Liu, Kevin Ritter, Verena B. Eisenbeis, Tamara Bittner, Artiom Gruzdev, Lea Seidel, Bertram Bengsch, Stephen B. Shears, Henning J. Jessen

## Abstract

Technical challenges have to date prevented a complete profiling of the levels of myo-inositol phosphates (InsPs) and pyrophosphates (PP-InsPs) in mammalian tissues. Here, we have deployed capillary electrophoresis mass spectrometry to identify and record the levels of InsPs and PP-InsPs in several tissues obtained from wild type mice and a newly-created PPIP5K2 knockout strain. We observe that the mouse colon harbours unusually high levels of InsPs and PP-InsPs. Additionally, the PP-InsP profile is considerably more complex than previously reported for animal cells: using chemically synthesized internal stable isotope references, and high-resolution mass spectra, we characterize two new PP-InsP isomers as 4/6-PP-InsP_5_ and 2-PP-InsP_5_. The latter has not previously been described in Nature. Analysis of feces and the commercial mouse diet suggest the latter is one potential source of noncanonical isomers in the colon. However, we also identify both molecules in the heart, indicating unknown synthesis pathways in mammals. We also demonstrate that the CE-MS method is sensitive enough to measure PP-InsPs from patient samples such as colon biopsies and peripheral blood mononuclear cells (PBMCs). Strikingly, PBMCs also contain 4/6-PP-InsP_5_ and 2-PP-InsP_5_. In summary, our study substantially expands PP-InsP biology in mammals.

---

The inositol phosphates (InsPs) and pyrophosphates (PP-InsPs) are a complex signalling hub with diverse functions in eukaryotes.^1-3^ The PP-InsPs have specialized physicochemical properties and biological functions that attract widespread interest.^4-7^ They occur as distinct isomers of differentially phosphorylated metabolites of InsP_6_ (phytic acid, phytate). The current literature suggests that in yeast and mammals these phosphorylation reactions occur selectively and successively in the 5- and 1-positions (Figure 1) leading to 5-PP-InsP_5_ and 1,5-(PP)_2_-InsP_4_, respectively.^8,9^ In plants and slime-mold, 4/6-PP-InsP_5_ has been identified as the main isomer, with the absolute configuration of the biologically relevant enantiomer remaining unknown.^10,11^

**Figure 1.**
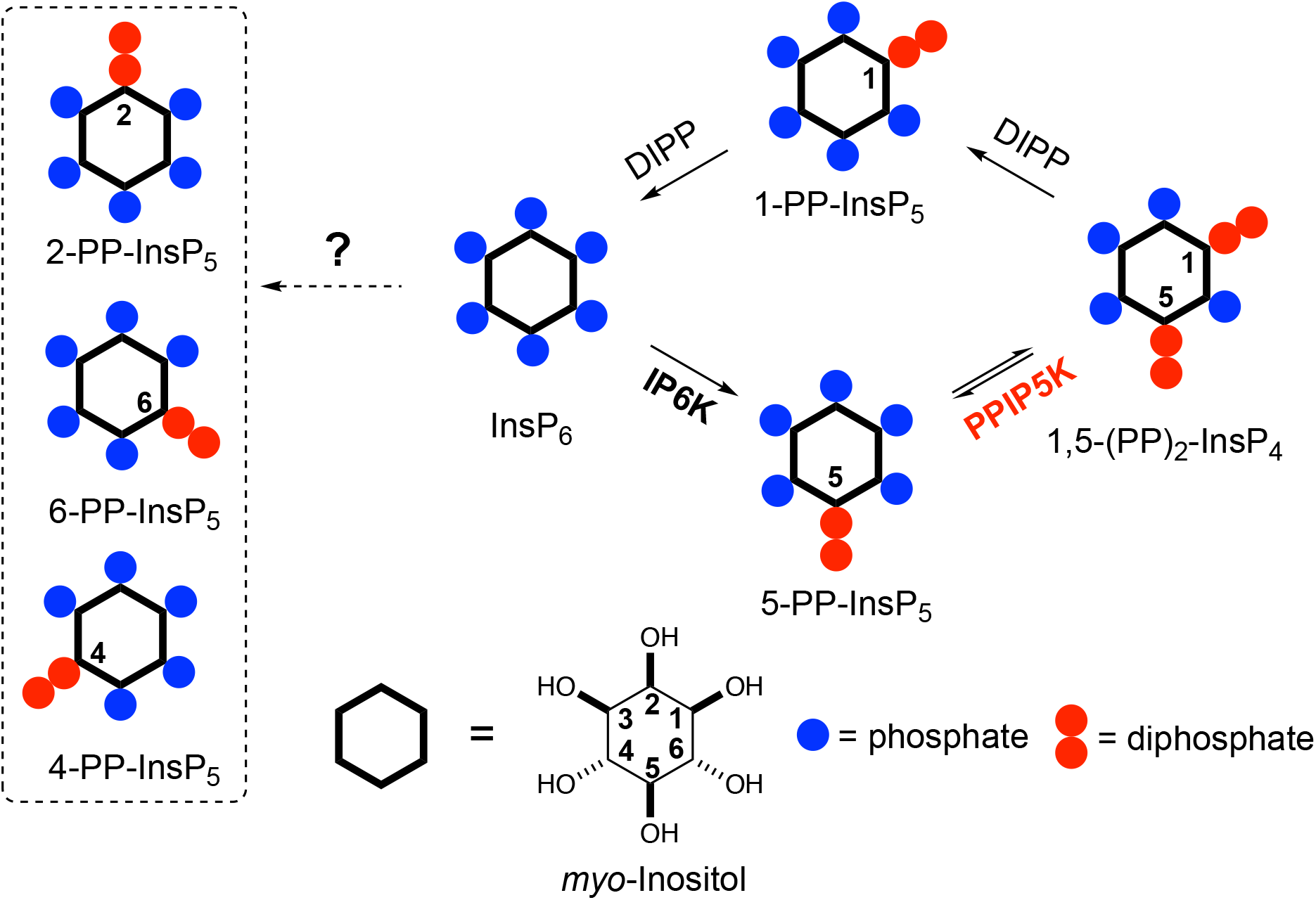
Main metabolic reactions that determine the turnover of inositol pyrophosphates in mammalian cells. Three isoforms of IP6Ks phosphorylate the 5-position of InsP_6_, two isoforms of PPIP5Ks phosphorylate the 1-position with a preference for 5-PP-InsP_5_ over InsP_6_. The question mark indicates that an unknown pathway is responsible for the synthesis of the 4/6-PP-InsP_5_ and 2-PP-InsP_5_ identified in the current study (inside dotted box, note that 4/6-PP-InsP_5_ are enantiomers). PPIP5Ks also harbour a phosphatase domain catalyzing dephosphorylation 1,5-(PP)_2_-InsP_4_. DIPPs are specialized phosphatases that degrade the phosphoric anhydrides in PP-InsPs.

Kinases and phosphatases that synthesize and metabolize PP-IPs are distributed throughout all eukaryotic kingdoms.^7,12^ In mammals, there are three isoforms of IP6Ks that add a β-phosphate at position 5, and two isoforms of PPIP5Ks that add a β-phosphate at position 1.^8^ Most of the research into PP-InsP turnover in mammalian cells has relied on separation by HPLC of extracts of ^3^H-inositol radiolabeled cells, although in more recent years a more generally-accessible PAGE technique has proved useful.^13,14^ This body of work has consistently concluded that 5-PP-InsP_5_ is the most abundant PP-InsP isomer (generally <10% of InsP_6_ levels). Levels of 1,5-(PP)_2_-InsP_4_ and 1-PP-InsP_5_ are approximately 10-fold and 50-fold lower, which are below the PAGE detection limit.^14-16^ The relative ease with which 5-PP-InsP_5_ abundance can be measured has in large part driven the field’s primary focus on this isomer. For example, this PP-InsP has been reported to regulate insulin signalling, exocytosis, processing body formation, intracellular protein localization, and bioenergetic homeostasis.^17-22^

More recently, 1,5-(PP)_2_-InsP_4_ has emerged as an independently-regulated cellular signal. This facet of PP-InsP signalling first arose from kinetic assessments^23^ of the PPIP5K kinase domain that phosphorylates 5-PP-InsP_5_ to 1,5-(PP)_2_-InsP_4_ and the separate phosphatase domain that degrades 1,5-(PP)_2_-InsP_4_ back to 5-PP-InsP_5_ (see Figure 1). Moreover, the phosphatase activity is inhibited by elevations in cellular levels of inorganic phosphate (P_i_), thereby enhancing net 1,5-(PP)_2_-InsP_4_ production independently of any changes in 5-InsP_7_ levels. ^23,24^ As a consequence, the net kinase and phosphatase activities are tied to cellular energy and phosphate homeostasis.^3,25^ It has since been demonstrated that 1,5-(PP)_2_-InsP_4_ stimulates P_i_ efflux from mammalian cells through an interaction with an SPX domain on the transmembrane XPR1 protein.^26,27^ Moreover, pharmacologic inhibition of IP6Ks in mammals (rodents and monkeys), which restrains PP-InsP_5_ and 1,5-(PP)_2_-InsP_4_ synthesis (see Figure 1), leads to attenuation of systemic hyperphosphatemia through inactivation of XPR1; these findings are an important milestone for potential pharmacological treatment of chronic kidney disease.^28^ Naturally-occurring human variants of PPIP5K2 have been associated with deafness^29^ and keratoconus.^30^ Recently, [^3^H]inositol-radiolabeling of a hematopoietic stem cell line from a PPIP5K2^-/+^ mouse indicated 1,5-(PP)_2_-[^3^H]InsP_4_ levels are no different from those in typical culture medium (data for PPIP5K2^-/-^ cells were not reported).^31^ In such circumstances, it has become more important to be able to accurately assay dynamic fluctuations in 1,5-(PP)_2_-InsP_4_ concentrations.

A portfolio of additional methods has been introduced that can assay mass levels of (PP)-InsPs in extracts of mammalian and plant cells, including using transition metals (e.g. Fe, Y) and absorbance detection (metal dye detection, MDD) ^32-34^ and the coupling of in-line mass spectrometry to hydrophilic interaction liquid chromatography (HILIC) and metal-free C_18_ reversed phase columns. ^28,35,36^ NMR detection with ^13^C enriched inositol is another recent and promising addition to the analytical portfolio.^37^ In 2020, capillary electrophoresis (CE) with mass spectrometry compatible buffers was reported for PP-InsP analytics, with only nanoliter sample consumption and accurate isomer assignment and quantitation by using stable isotope internal reference compounds. ^38^

We now significantly expand the value of our new PP-InsP profiling techniques through our identification of substantial cellular quantities of mammalian 4/6-PP-InsP_5_ and 2-PP-InsP_5_ (see Figure 1) based on comigration with reference compounds and high-resolution mass spectra. This conclusion is facilitated by adapting a recently developed ^18^O phosphate labelling approach^39^ in order to stereoselectively synthesize 4-PP-InsP_5_ to use as a heavy internal standard. Finally, it was our goal to optimize CE-MS to monitor the complete array of PP-InsPs from human patient tissues. For this work, we selected colon biopsies and peripheral blood mononuclear cells including enriched T cell subpopulations (PBMCs, CD8^+^). Strikingly, we also identify 4/6-PP-InsP_5_ and 2-PP-InsP_5_ in PBMCs that are particularly enriched in a CD8^+^ T cell preparation. Overall, this dramatic increase in the complexity of PP-InsP metabolism indicates that their biological significance has been greatly underestimated.

## Results

With an established protocol that uses TiO_2_ beads, we extracted and enriched InsPs and PP-InsPs from different mouse tissues. ^14,38^ The enriched samples were analyzed by CE-QQQ using the same background electrolyte (35mM ammonium acetate adjusted to pH 9.7 with NH_4_OH, i.e., BGE-A) that we deployed in our previous study.^38^ Samples were spiked with internal heavy isotope reference compounds (^13^C labels) of several different InsPs and PP-InsPs for assignment and quantitation. This is the first time this method has been applied to any animal tissue for the quantification of the levels of the least abundant PP-IPs, namely 1,5-(PP)_2_-InsP_4_ and 1-PP-InsP_5_ (for representative examples see Figure 2C and Supplementary Figure 1).

**Figure 2.**
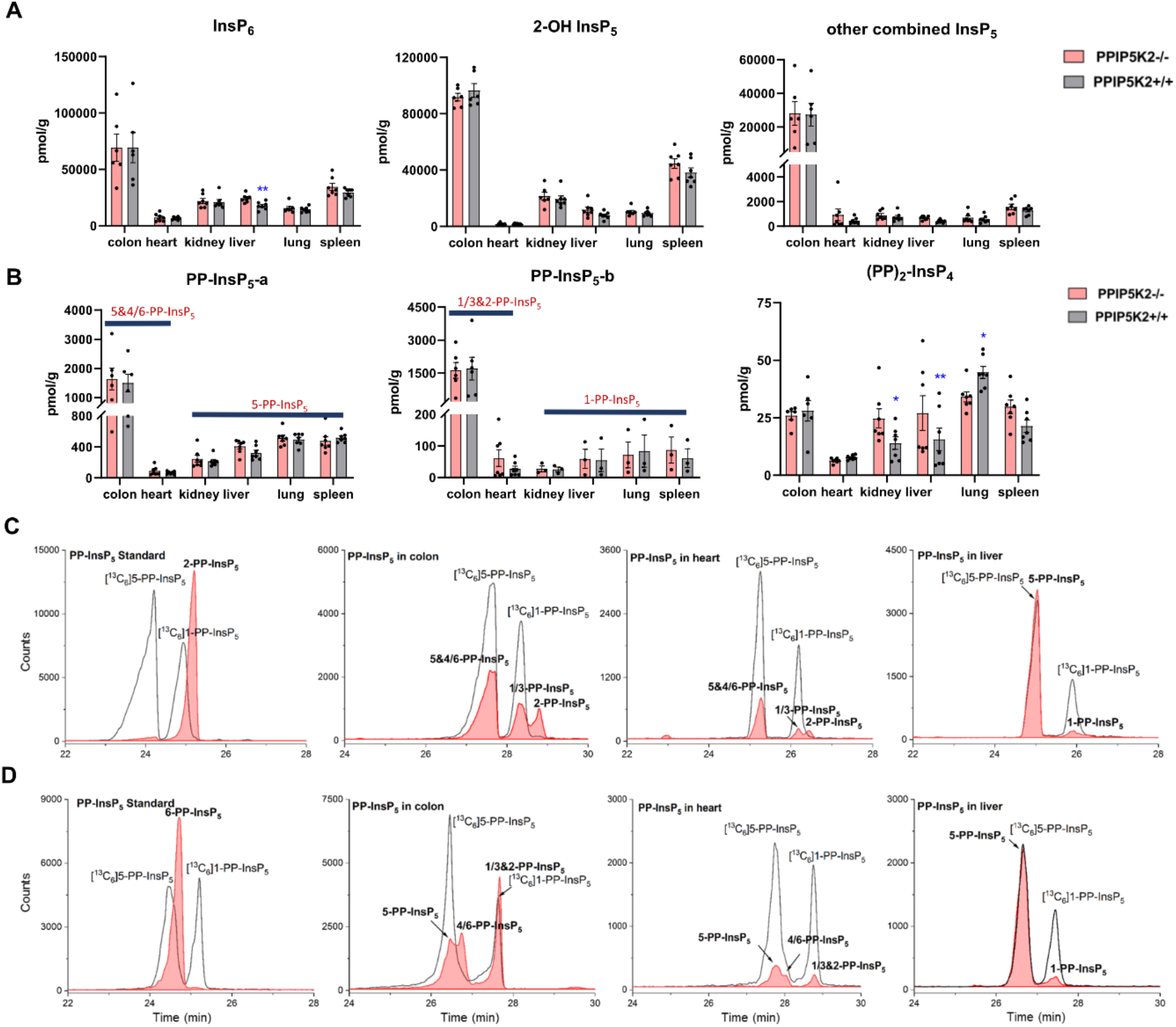
Profiling of PP-InsPs in mouse tissues (Wild type vs PPIP5K2 knockout) and observation of new isomers. **A** Profiling of InsP_6_, 2-OH InsP_5_, and the total for all other InsP_5_ isomers (4/6-OH InsP_5_, 1/3-OH InsP_5_, 5-OH InsP_5_). **B** PP-InsP_5_-a and PP-InsP_5_-b refer to two base-line resolved peaks. The tentative identification of the components of each peak (in bold font) is described in the text. Data for A and B are indicated as means +/- SEM, n=7 and for colon, n=6. *P < 0.05 and **P < 0.01, Student’s t-test. **C** Extracted ion electropherograms (EIEs) of [_13_C] labeled internal reference compounds (black lines) plus a 2-PP-InsP_5_ standard (red trace) or PP-InsP_5_ extracts from mouse colon, heart and liver, resolved with BGE A containing 35 mM ammonium acetate titrated with ammonium hydroxide to pH 9.7. **D** EIEs of [^13^C]-labeled internal reference compounds (black lines) plus (left panel) 6-PP-InsP_5_ standard (red trace) or PP-InsP_5_ extracts from mouse colon, heart and liver (red trace), resolved with BGE B containing 40 mM ammonium acetate titrated with ammonium hydroxide to pH 9.0.

We also used this method to compare InsP and PP-InsP levels in multiple mouse tissues, including colon, heart, kidney, liver, lung and spleen (Figure 2A,B,C). These molecules were generally least abundant in the heart. It is worth mentioning that other minor InsP_5_ isomers including 4/6-OH InsP_5_, 1/3-OH InsP_5_, 5-OH InsP_5_ have also been identified and quantified (see representative examples obtained from mouse colon and mouse heart; Supplementary Figure 2), while 2-OH InsP_5_ was always by far the predominant isomer in all investigated mouse tissues (Figure 2A).

Compared to other tissues, the colon is notable for containing substantially higher levels of InsP_6_ (2- to 5-fold), 2-OH-InsP_5_ (2- to 10-fold) and the sum of the remaining, quantitatively more minor InsP_5_ isomers (19- to 52-fold). The colon also contains much higher levels of PP-InsP_5_ isomers (Figure 2A,B,C). In most of the studied tissues (kidney, liver, lung, and spleen), two baseline-resolved PP-InsP_5_ signals were observed (labeled ‘a’ and ‘b’), which co-eluted precisely with internal standards of [^13^C_6_]5-PP-InsP_5_ and [^13^C_6_]1-PP-InsP_5_, respectively, in each of two different BGE conditions (Figure 2C, Supplementary Figure 3A). In these tissues, the relative proportion of 1-PP-InsP_5_ to 5-PP-InsP_5_ (approximately 1 to 7) is higher than that determined by our previous CE analysis of a line of immortalized HCT116 cells (1 to 13)^40^; a ratio of only 1 to 50 was previously obtained by HPLC analysis of [^3^H]inositol-labeled extracts of immortalized cells.^41^

An unexpected outcome of the EIE obtained using BGE-A was that the PP-InsP-b signals derived from the colon and heart split into two approximately equally-sized peaks that are incompletely resolved; the earlier-eluting peak comigrated with an internal standard of [^13^C_6_]1-PP-InsP_5_ (Figure 2C). The elution time of the second peak corresponds precisely to the elution time of a replicate sample spiked with an internal standard of 2-PP-InsP_5_ (Supplementary Figure 4). In addition, there is an indication that the PP-InsP-a signal derived from the colon also separates into two incompletely resolved peaks (Figure 2C). To pursue the latter observation, we reran the samples with the background electrolyte adjusted to 40 mM ammonium acetate titrated with ammonium hydroxide to pH 9.0 (i.e., BGE-B). This procedure extended the peak-to-peak resolution within the PP-InsP-a signal to the extent that its two components are also visible in the extracts prepared from the colon and heart (Figure 2D). Note that, in contrast, the use of BGE-B did not perturb the coelution of internal standards of [^13^C_6_]5-PP-InsP_5_ and [^13^C_6_]1-PP-InsP_5_ with PP-InsP-a and PP-InsP-b signals, respectively, that were prepared from kidney, liver, lung and spleen (Figure 2C,D; Supplementary Figure 3A,B). However, we do not exclude that matrix effects in other tissues would blur the presence of low levels of additional PP-InsP isomers.

In this set of experiments with BGE-B, the first component of PP-InsP-a extracted from the colon comigrates with the internal standard of [^13^C_6_]5-PP-InsP_5_ and the second component of PP-InsP-a has an elution time that matches that of a standard of 6-PP-InsP_5_ from separate runs (Figure 2D). Thus, we tentatively identify the second component of PP-InsP-a as 4/6-PP-InsP_5_ and by a process of elimination we suggest the second component of PP-InsP-b is 2-PP-InsP_5_. Moreover, the proposed nature of 1/3-PP-InsP_5_, 2-PP-InsP_5_, 5-PP-InsP_5_ and 4/6-PP-InsP_5_ from the colon is also consistent with their high-resolution mass spectra collected by a CE-qTOF system (Supplementary Figure 5). Other potential candidates with an identical mass, such as triphosphates of inositol-tetrakisphosphates (e.g. 5-PPP-InsP_4_), have been described so far only *in vitro*.^42^ The *myo*-configuration for these new PP-InsPs seems likely, since there is no prior identification of any other multiply phosphorylated inositol stereoisomers in mammals.

It is notable that in the colon we estimate the levels of 1-PP-InsP_5_ (i.e., half of PP-InsP-b) and 5-PP-InsP_5_ (i.e., half of PP-InsP-a) are approximately equivalent (Figure 2C,D); this observation implies we must profoundly modify prior perceptions of 1-PP-InsP_5_ as a quantitatively minor constituent of mammalian cells, and/or consider the possibility that the enantiomer 3-PP-InsP_5_ is also present. Currently applied methods do not resolve the enantiomers.

The 2-PP-InsP_5_ isomer has not previously been identified in any biological material, possibly because it is both unexpected and only present at relatively low levels. On the contrary, 4/6-PP-InsP_5_ was recently discovered to be a major PP-InsP isomer in plants. ^10^ Clearly, the latter is also a quantitatively important isomer in the mouse colon and heart (Figure 2), and so it was particularly important to further validate its nature. Thus, we have developed a synthetic route to the preparation of enantiomerically pure [^18^O_2_]4-PP-InsP_5_ to deploy as an internal standard for additional chromatographic resolutions (see Figure 3).

**Figure 3.**
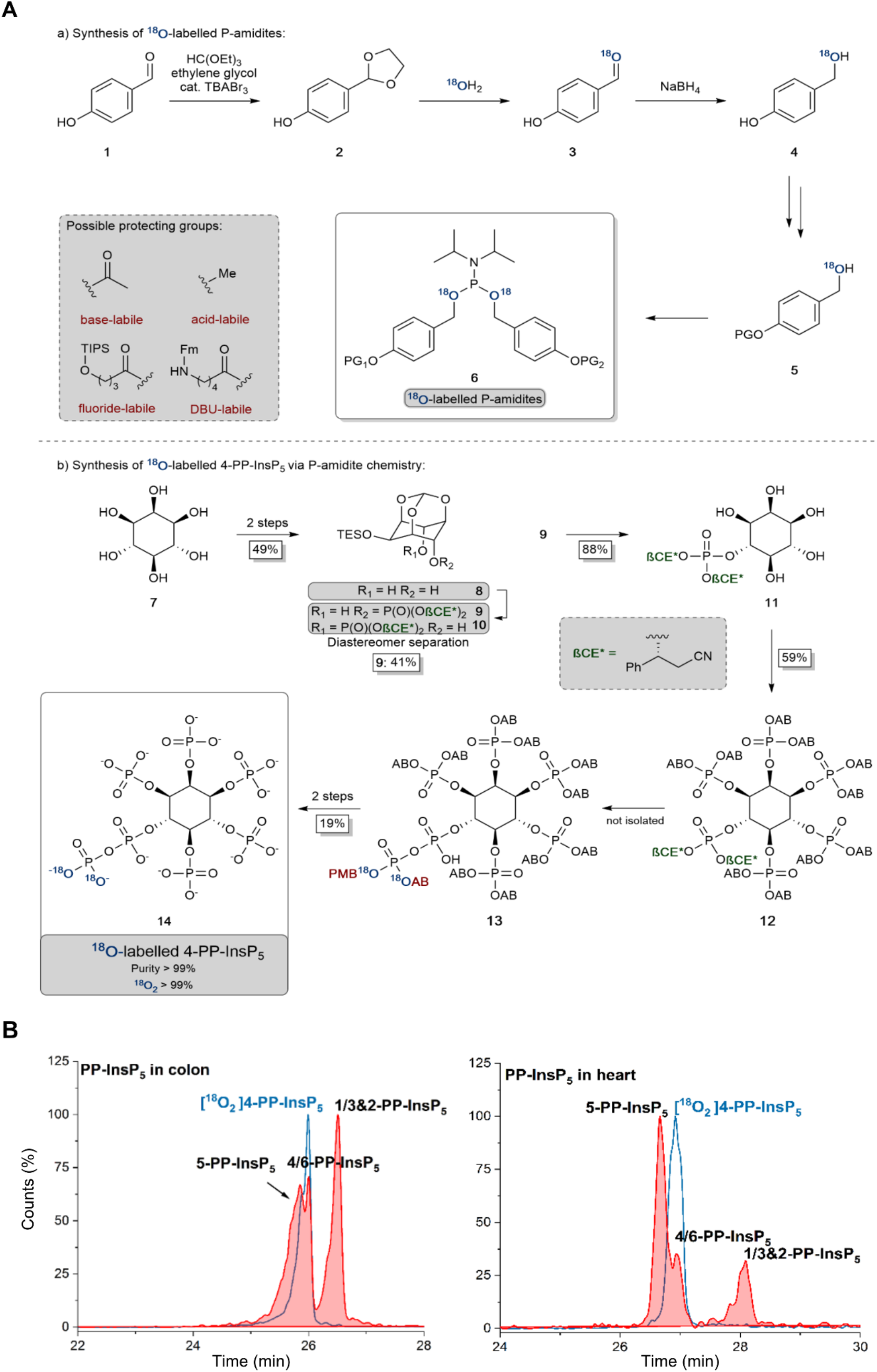
**A:** Synthesis and application of _18_O labeled P-amidites with diverse protecting group patterns and their application to a late-stage labeling 4-PP-InsP_5_ synthesis. AB: Acetoxybenzyl, PMB: para-methoxybenzyl **B:** Separation of 5-PP-InsP_5_ and 4/6-PP-InsP_5_ (filled red plots) from mouse colon and heart samples using BGE-B and assignment of the isomer with the new internal reference compound [^18^O_2_] 4-PP-InsP_5_ (blue plot) as either 4-PP-InsP_5_ or 6-PP-InsP_5_. EIEs (PP-InsP_5_ and [^18^O_2_] PP-InsP_5_) are scaled to the largest peak indicated as 100%.

We have also recorded 1,5-(PP)_2_-InsP_4_ levels in mouse tissues (Figure 2B). These varied over a 5-fold range, with the lowest levels in the heart and the highest in the lung; as far as we are aware, no previous study has provided such data. This accomplishment enabled us to determine the impact upon 1,5-(PP)_2_-InsP_4_ levels in a newly-created PPIP5K2 knockout mouse (Supplementary Figure 6). The knockout only resulted in a statistically significant reduction in 1,5-(PP)_2_-InsP_4_ levels in the lung tissue (Figure 2B). In fact, 1,5-(PP)_2_-InsP_4_ levels trended higher in several PPIP5K2 knockout tissues compared to the wild-type, and in kidney and liver this effect was statistically significant. Although this might initially seem a counter-intuitive outcome, it is possible that in these two tissues loss of the PPIP5K2 1,5-(PP)_2_-InsP_4_-phosphatase domain may have a larger metabolic effect than loss of the 5-PP-InsP_5_ kinase domain. The knockout did not elicit a statistically-significant impact on 1,5-(PP)_2_-InsP_4_ levels in either the colon, or heart. The observation of tissue dependent variability in PP-InsP signaling brought about by PPIP5K2 knockout may depend in part due to the extent to which PPIP5K1 compensates for deletion of PPIP5K2 catalytic activity, although no such effect was evident in liver (Supplementary Figure 6). Note also that the PPIP5K2 KO did not have off-target effects on any of the other InsPs and PP-InsPs analyzed in this study (Figure 2A,B), except that InsP_6_ was increased in the PPIP5K KO liver.

We could not derive sufficient purified amounts of the putative 4/6-PP-InsP_5_ for NMR analysis to further corroborate the identity of this isomer. So instead, we generated a reference compound with a heavy isotope label to serve as an internal standard for CE-MS. We reasoned that comigration of this compound under different separation conditions would serve as a strong indication that it is indeed 4/6-PP-InsP_5_ in its *myo*-configuration. The enzymes for plant 4/6-PP-InsP_5_ synthesis are not yet known and so an enzymatic synthesis starting from InsP_6_ of the reference compound with ^13^C labels was not possible.^43^ A fully chemical synthesis from expensive ^13^C glucose in a multi-step linear approach was deemed not feasible.^37^ We thus relied on our recently developed ^18^O phosphate labeling approach in which the expensive isotopic label can be introduced in the penultimate step of the synthesis.^39^

In brief, ^18^O labeled phosphoramidites (P-amidites) with high ^18^O/^16^O ratios are key to the synthesis. These high ratios can be obtained by the strategy shown in figure 3A a). Para-hydroxybenzaldehyde is transformed into its acetal **2**, which is then hydrolyzed in the presence of 99% ^18^O enriched water. The aldehyde **3** is directly reduced to stable alcohol **4**, which can then be protected on the phenol with diverse protecting groups (in the case described here simply acetate giving the acetoxybenzyl (AB) protecting group). The alcohols **5** are then transformed into P-amidites of the general structure **6**, enabling diverse protecting group patterns and high ^18^O enrichment. The inositol structure is assembled as reported previously,^44-47^ shown in figure 3A b). While strictly a desymmetrization was not required and the generation of racemic 4/6-PP-InsP_5_ would have been sufficient, we still generated the enantiomerically pure compound for potential future applications. Desymmetrization was achieved from intermediate protected diol **8**, which was reacted with an unsymmetric P-amidite containing chiral protecting groups (β-CE*, an arylated enantiomerically pure variant of the β-cyanoethyl protecting group). The obtained diastereomeric mixture was separated and then the inositol protecting groups were removed giving pentaol **11. 11** was phosphorylated to protected InsP_6_ **12** with orthogonal protecting groups (β-CE*) in the 4-position.^44^ Selective deprotection in that position then enables the introduction of the ^18^O labeled phosphate bearing two ^18^O oxygen atoms (M+4). Global deprotection gave [^18^O]_2_ 4-PP-InsP_5_ **14** in 99% purity with >99% isotopic enrichment as determined by CE-MS. This reference compound was then dissolved in water and its concentration was determined by quantitative ^1^H- and ^31^P-NMR.

Figure 3B demonstrates the first application of this newly generated isotopologue. Briefly, both colon and heart samples were spiked with the new reference and we utilized the optimized BGE-B that is capable of 5-PP-InsP_5_ and 4/6-PP-InsP_5_ separation. Masses were recorded and an identical migration of the unknown analyte with our reference in the same matrix was found, strongly suggesting it is indeed 4/6-PP-InsP_5_ that has been measured for the first time in mammalian tissues.

To understand the complexity of the profiles of InsPs and PP-InsPs in the colon in an organismal context, we additionally analyzed mouse feces and found them to contain very high levels of most analytes (Figure 4A and C, Supplementary Figure 7). Moreover, neither PP-InsP peak co-eluted precisely with internal standards of either 5-PP-InsP_5_ or 1-PP-InsP_5_ again pointing towards the existence of both 4/6-PP-InsP_5_ and 2-PP-InsP_5_. In fact, the two new isomers are the most abundant analytes we detect (Figure 4A). Interestingly, the PPIP5K knockout contained increased levels of all analytes in feces. We excluded that the colon microbiome contributed to the PP-InsP profile, because a comparison of germ-free and regular mouse colon samples did not show large changes in either identity or abundance (Supplementary Figure 8).

**Figure 4.**
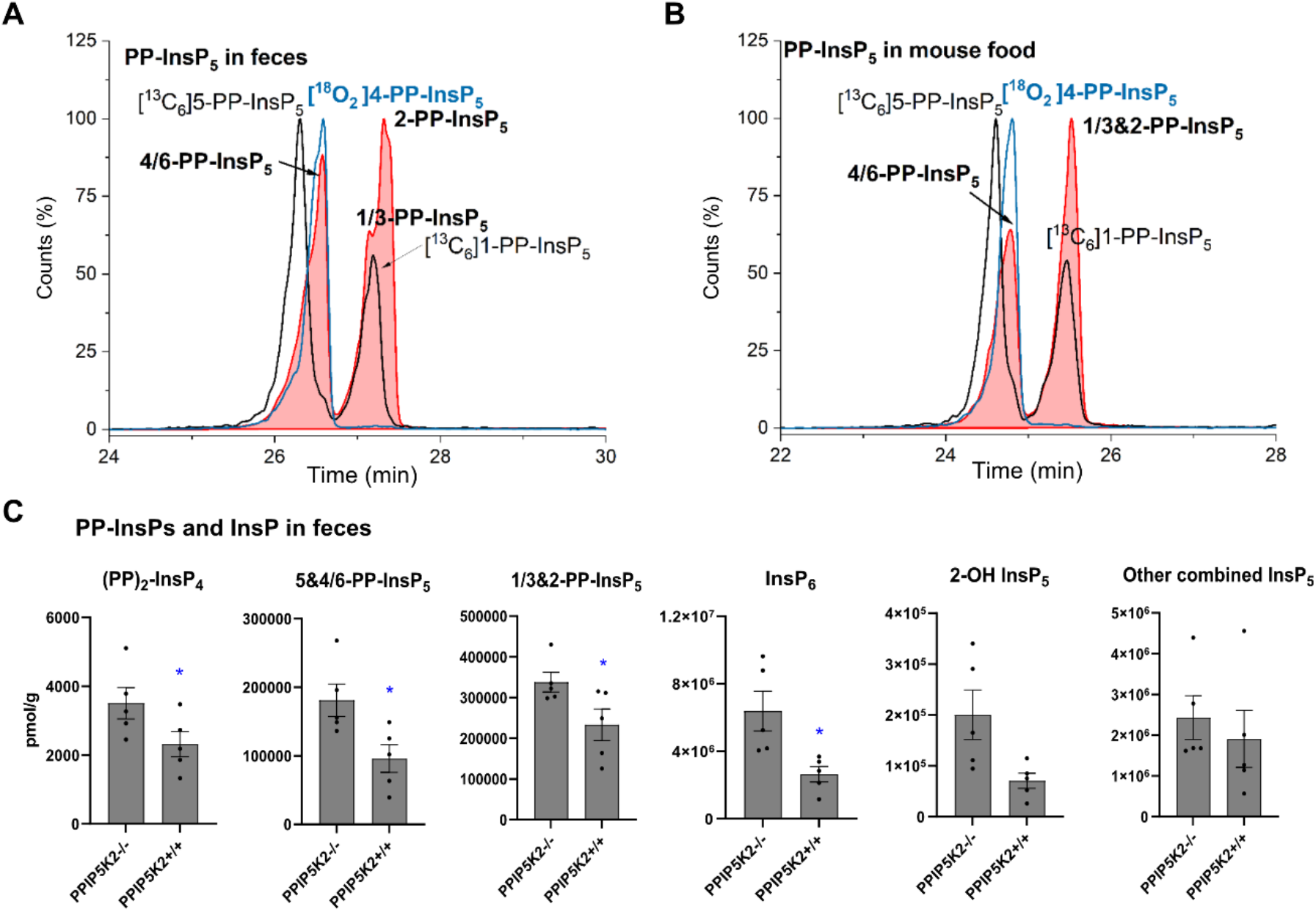
**A** CE-MS analysis of PP-InsP_5_ in mouse feces (filled red plots) with internal references of [^13^C_6_]5-PP-InsP_5_ and [^13^C_6_]1-PP-InsP_5_ (black plot) and [^18^O_2_]4-PP-InsP_5_ (blue plot) using BGE B. 4/6-PP-InsP_5_ isomer is identified in mouse feces as well. **B** Analysis of PP-InsP_5_ in mouse food same as in A, which contains high levels of this 4/6-PP-InsP_5_ isomer. **C** Profiling of PP-InsPs and InsPs in mouse feces in (Wild type vs PPIP5K2 knockout). Data are indicated as means +/- SEM, n=5. *P < 0.05, Student’s t-test.

We next investigated if the mouse laboratory diet might contribute to the unprecedented complexity of the colonic PP-InsP profile. We provided mice with the “Rodent NIH-31 Open Formula Autoclavable Diet”, much of which is of plant origin. This is significant because recent work has determined that the quantitatively most important PP-InsP isomer in plants is one that had previously been overlooked, namely, 4/6-PP-InsP_5._^10^ Indeed, our internal standards allowed us to conclude that large amounts of 4/6-PP-InsP_5_ were present in the mouse diet, although a precise quantification was hindered by insufficient separation of the 4/6- and 5-PP-InsP_5_ peaks from within the PP-InsP_5_-a peak (Figure 4**B**). Nevertheless, the latter was smaller than the PP-InsP_5_-b peak, which likely comprises a mixture of 1-PP-InsP_5_ and 2-PP-InsP_5_. These data imply plants utilize a pathway to 2-PP-InsP_5_ synthesis.

Our results also raise the possibility that the diet might be the source of the colon’s unusually high levels of InsP_6_ and PP-InsPs, as well as the more complex PP-InsP profile. Furthermore, 2-OH InsP_5_ is the minor InsP_5_ isomer in mouse feces and also in mouse food (Supplementary Figure 9**A**), in contrast to it being the major InsP_5_ in the colon. This result suggests that the exceptional PP-InsPs and InsP_6_ profile in colon are not due to contamination from feces during sample preparation. In this case, endocytosis of dietary InsP_6_ and PP-InsPs by colonic epithelial cells should be considered as a viable possibility.

Finally, in order to demonstrate the sensitivity of the method and its potential in translational research, we obtained human samples for enrichment and profiling. We analyzed one 18 mg wet tissue colon biopsy, which was sufficient to profile the main PP-InsP and InsP content (Figure 5A). Only canonical isomers were identified, i.e. 5-PP-InsP_5_, InsP_6_, and 2-OH InsP_5_. We additionally analyzed peripheral blood mononuclear cells (PBMCs; Figure 5B) from donors, and also CD8^+^ T-cells enriched from the PBMC pool by FACS (see Supplementary Information). Strikingly, in one such enriched sample, we identified 4/6-PP-InsP_5_ as the sole PP-InsP isomer (Figure 5C). Of note, the CD8^+^ depleted PBMC pool (Figure 5D) also contained 4/6-PP-InsP_5_ as well as 5-PP-InsP_5_ and the latter was identified as the minor isomer. Moreover, a peak comigrating with 2-PP-InsP_5_ was identified in PBMCs (Figure 5B) and can be tentatively assigned in a shoulder of the peak of the CD8^+^ depleted fraction (Figure 5D). CD8^+^ enrichment did not provide enough material for analysis in all samples studied, so it remains unclear whether the surprising 4/6-PP-InsP_5_ enrichment is generally found in CD8^+^ cells from different donors. However, our analysis now firmly establishes this new isomer is of mammalian origin.

**Figure 5.**
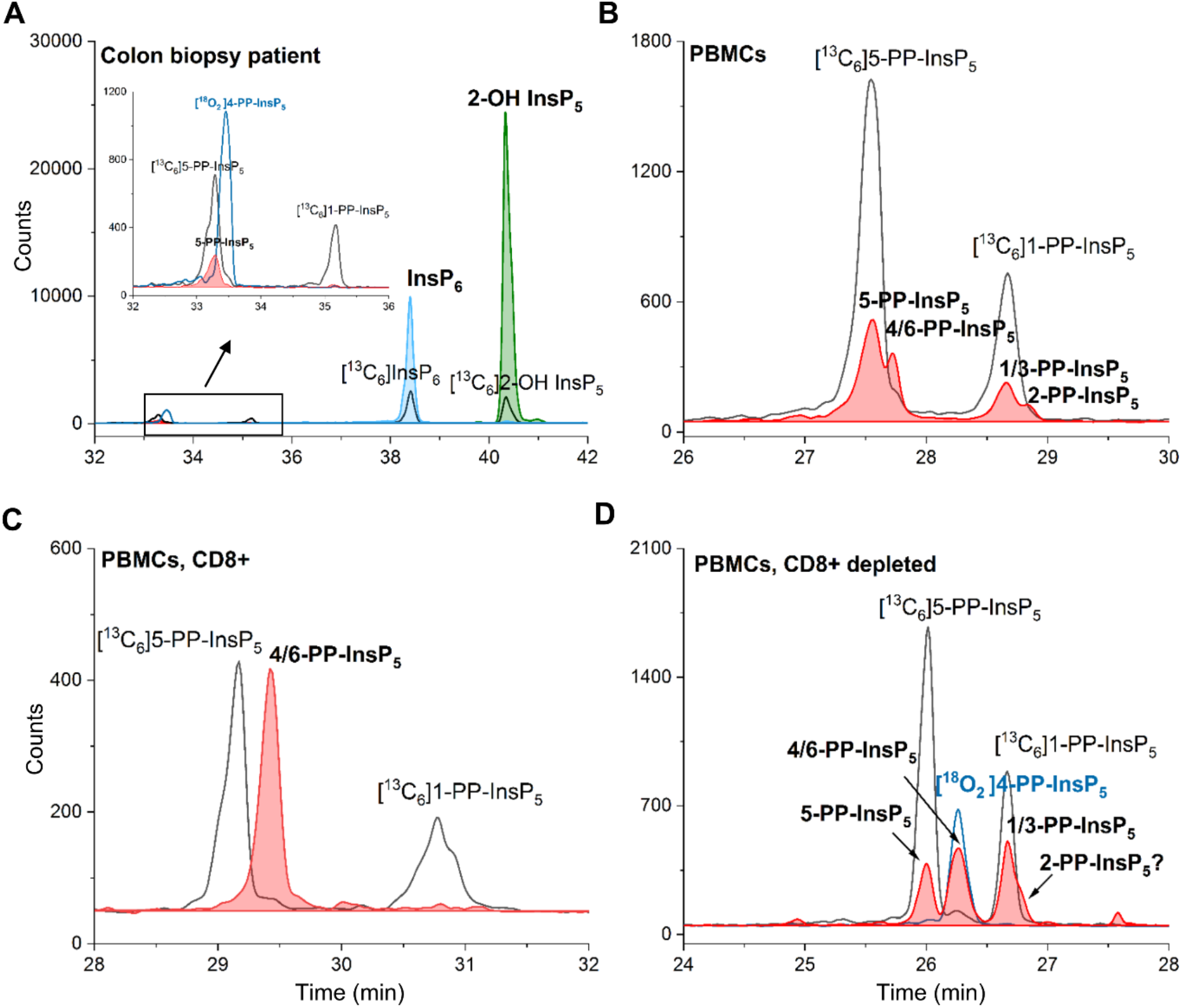
**A** CE-MS Analysis of a human colon tissue biopsy (18 mg) enables identification of several important inositol phosphate (InsP_6_ and InsP_5_) and pyrophosphate isomers (5-PP-InsP_5_). **B** 4/6-PP-InsP_5_ is identified in PBMCs by CE-MS analysis. The electropherograms are representative of independent biological triplicates giving comparable results. **C** 4/6-PP-InsP_5_ is enriched in a CD8^+^ T-cell preparation and is also present in the CD8_+_ depleted PBMC pool (**D**). It is assigned by its exact same migration time as [^18^O_2_] 4-PP-InsP_5_. PP-InsP_5_ (filled red plot) isomer identification is achieved with the aid of [^13^C_6_]5-PP-InsP_5_, [^13^C_6_]1-PP-InsP_5_ (black plot) and [^18^O_2_] 4-PP-InsP_5_ (blue plot).

## Conclusions

We have applied CE-MS profiling to delineate a more sophisticated picture of InsP and PP-InsP distributions in metazoan samples. Thus, inositol pyrophosphate signalling appears even more complex than previously thought. The CE-MS method also has sufficient sensitivity to profile for the first-time biopsies from human patients and PBMCs including isolated CD8^+^ T-cells from human blood. We obtain several unexpected results based on the high separation efficiency of capillary electrophoresis that have gone undetected with recently developed LC-MS approaches.^28,35,36^ In particular, we identify very high levels of PP-InsPs in colon tissue, which are potentially endocytosed from the laboratory diet, including large quantities of the putative noncanonical 4/6 and 2-PP-InsP_5_ isomers. Our data therefore represent a paradigm shift in our understanding of dietary influences upon PP-InsP metabolism and signaling in the colon. While 4/6-PP-InsP_5_ and 2-PP-InsP_5_ in colon could possibly originate from endocytosis of food constituents, this phenomenon cannot apply to heart samples as well as human PBMCs. Consequently, it appears that 4/6-PP-InsP_5_ and 2-PP-InsP_5_ can also be synthesized by mammals.

Our new isomer assignments are based on the exact mass determination and exact comigration with standards of both PP-InsPs, including a novel synthetic 4-PP-InsP_5_ bearing two ^18^O oxygen isotope labels. Future studies must now address the enantiomeric identity of the new metazoan 4/6-PP-InsP_5_ as well as a complete structural assignment of 2-PP-InsP_5_ by NMR to firmly establish *myo*-configuration and exclude other potential constitutional isomers of the same mass and identical migration during CE. The colonic uptake, dynamic regulation, unknown enzymology and functions of these new isomers will be productive directions for future research. With the ability to profile PP-InsPs from human biopsies and blood samples, their establishment as potential disease biomarkers will also become an important future endeavor.

## Author contributions

DQ and CG designed, performed, and evaluated analytical experiments. CG was responsible for the animal experiments. SBS supervised animal experiments. GL and VE performed biopsy extractions and analytics. KR conducted the chemical synthesis. TB and LS isolated and extracted PBMCs. BB supervised human tissue extractions and designed experiment. HJJ and SBS conceived of the project and designed experiments. HJJ, DQ, CG, SBS wrote the manuscript. All authors provided feedback on experimental design and contributed to manuscript revisions.

## Acknowledgements

This study was supported by the German Research Foundation (DFG) under Germany’
ss excellence strategy (CIBSS – EXC-2189 – Project ID 390939984) and DFG Grant JE 572/4-1. HJJ and GL acknowledge funding from the Volkswagen Foundation (VW Momentum Grant 98604). This research was also supported by the Intramural Research Program of the NIH, National Institute of Environmental Health Sciences. The authors wish to thank Dorothea Fiedler (Leibniz Institut für Molekulare Pharmakologie, Berlin) and particularly Minh Nguyen Trung und Robert Harmel for providing ^13^C-labeled reference compounds.

## References

(1) Irvine, R. F.; Schell, M. J. Back in the water: The return of the inositol phosphates. Nat Rev Mol Cell Bio 2001, 2, 327.

(2) Laha, D.; Portela-Torres, P.; Desfougeres, Y.; Saiardi, A. Inositol phosphate kinases in the eukaryote landscape. Adv Biol Regul 2021, 79, 100782.

(3) Shears, S. B. Intimate connections: Inositol pyrophosphates at the interface of metabolic regulation and cell signaling. J Cell Physiol 2018, 233, 1897.

(4) Lee, S.; Kim, M. G.; Ahn, H.; Kim, S. Inositol Pyrophosphates: Signaling Molecules with Pleiotropic Actions in Mammals. Molecules 2020, 25.

(5) Shears, S. B.; Wang, H. Metabolism and Functions of Inositol Pyrophosphates: Insights Gained from the Application of Synthetic Analogues. Molecules 2020, 25.

(6) Brown, N. W.; Marmelstein, A. M.; Fiedler, D. Chemical tools for interrogating inositol pyrophosphate structure and function. Chem Soc Rev 2016, 45, 6311.

(7) Nguyen Trung, M.; Furkert, D.; Fiedler, D. Versatile signaling mechanisms of inositol pyrophosphates. Current opinion in chemical biology 2022, 70, 102177.

(8) Shears, S. B. Inositol pyrophosphates: why so many phosphates? Adv Biol Regul 2015, 57, 203.

(9) Randall, T. A.; Gu, C.; Li, X.; Wang, H.; Shears, S. B. A two-way switch for inositol pyrophosphate signaling: Evolutionary history and biological significance of a unique, bifunctional kinase/phosphatase. Adv Biol Regul 2020, 75, 100674.

(10) Riemer, E.; Qiu, D.; Laha, D.; Harmel, R. K.; Gaugler, P.; Gaugler, V.; Frei, M.; Hajirezaei, M.-R.; Laha, N. P.; Krusenbaum, L.; Schneider, R.; Saiardi, A.; Fiedler, D.; Jessen, H. J.; Schaaf, G.; Giehl, R. F. H. ITPK1 is an InsP6/ADP phosphotransferase that controls phosphate signaling in Arabidopsis. Mol Plant 2021, 14, 1864.

(11) Desfougères, Y.; Portela-Torres, P.; Qiu, D.; Livermore, T. M.; Harmel, R. K.; Borghi, F.; Jessen, H. J.; Fiedler, D.; Saiardi, A. The inositol pyrophosphate metabolism of Dictyostelium discoideum does not regulate inorganic polyphosphate (polyP) synthesis. Adv Biol Regul 2022, 83, 100835.

(12) Kilari, R. S.; Weaver, J. D.; Shears, S. B.; Safrany, S. T. Understanding inositol pyrophosphate metabolism and function: kinetic characterization of the DIPPs. FEBS Lett 2013, 587, 3464.

(13) Losito, O.; Szijgyarto, Z.; Resnick, A. C.; Saiardi, A. Inositol Pyrophosphates and Their Unique Metabolic Complexity: Analysis by Gel Electrophoresis. Plos One 2009, 4.

(14) Wilson, M. S. C.; Bulley, S. J.; Pisani, F.; Irvine, R. F.; Saiardi, A. A novel method for the purification of inositol phosphates from biological samples reveals that no phytate is present in human plasma or urine. Open Biol 2015, 5, 150014.

(15) Pisani, F.; Livermore, T.; Rose, G.; Chubb, J. R.; Gaspari, M.; Saiardi, A. Analysis of Dictyostelium discoideum Inositol Pyrophosphate Metabolism by Gel Electrophoresis. Plos One 2014, 9, e85533.

(16) Pavlovic, I.; Thakor, D. T.; Bigler, L.; Wilson, M. S.; Laha, D.; Schaaf, G.; Saiardi, A.; Jessen, H. J. Prometabolites of 5-Diphospho-myo-inositol Pentakisphosphate. Angew Chem Int Ed 2015, 54, 9622.

(17) Chakraborty, A.; Koldobskiy, M. A.; Bello, N. T.; Maxwell, M.; Potter, J. J.; Juluri, K. R.; Maag, D.; Kim, S.; Huang, A. S.; Dailey, M. J.; Saleh, M.; Snowman, A. M.; Moran, T. H.; Mezey, E.; Snyder, S. H. Inositol Pyrophosphates Inhibit Akt Signaling, Thereby Regulating Insulin Sensitivity and Weight Gain. Cell 2010, 143, 897.

(18) Lee, T.-S.; Lee, J.-Y.; Kyung, J. W.; Yang, Y.; Park, S. J.; Lee, S.; Pavlovic, I.; Kong, B.; Jho, Y. S.; Jessen, H. J.; Kweon, D.-H.; Shin, Y.-K.; Kim, S. H.; Yoon, T.-Y.; Kim, S. Inositol pyrophosphates inhibit synaptotagmin-dependent exocytosis. Proc Natl Acad Sci USA 2016, 113, 8314.

(19) Sahu, S.; Wang, Z.; Jiao, X.; Gu, C.; Jork, N.; Wittwer, C.; Li, X.; Hostachy, S.; Fiedler, D.; Wang, H.; Jessen, H. J.; Kiledjian, M.; Shears, S. B. InsP<sub>7</sub> is a small-molecule regulator of NUDT3-mediated mRNA decapping and processing-body dynamics. Proc Natl Acad Sci USA 2020, 117, 19245.

(20) Shah, A.; Bhandari, R. IP6K1 upregulates the formation of processing bodies by influencing protein-protein interactions on the mRNA cap. J Cell Sci 2021, 134, jcs259117.

(21) Pavlovic, I.; Thakor, D. T.; Vargas, J. R.; McKinlay, C. J.; Hauke, S.; Anstaett, P.; Camuña, R. C.; Bigler, L.; Gasser, G.; Schultz, C.; Wender, P. A.; Jessen, H. J. Cellular delivery and photochemical release of a caged inositol-pyrophosphate induces PH-domain translocation in cellulo. Nat Commun 2016, 7, 10622.

(22) Szijgyarto, Z.; Garedew, A.; Azevedo, C.; Saiardi, A. Influence of Inositol Pyrophosphates on Cellular Energy Dynamics. Science 2011, 334, 802.

(23) Gu, C.; Nguyen, H.-N.; Hofer, A.; Jessen, H. J.; Dai, X.; Wang, H.; Shears, S. B. The Significance of the Bifunctional Kinase/Phosphatase Activities of Diphosphoinositol Pentakisphosphate Kinases (PPIP5Ks) for Coupling Inositol Pyrophosphate Cell Signaling to Cellular Phosphate Homeostasis. J Biol Chem 2017, 292, 4544.

(24) Dollins, D. E.; Bai, W.; Fridy Peter, C.; Otto James, C.; Neubauer Julie, L.; Gattis Samuel, G.; Mehta Kavi, P. M.; York John, D. Vip1 is a kinase and pyrophosphatase switch that regulates inositol diphosphate signaling. Proc Natl Acad Sci USA 2020, 117, 9356.

(25) Gu, C.; Nguyen, H.-N.; Ganini, D.; Chen, Z.; Jessen Henning, J.; Gu, Z.; Wang, H.; Shears Stephen, B. KO of 5-InsP7 kinase activity transforms the HCT116 colon cancer cell line into a hypermetabolic, growth-inhibited phenotype. Proc Natl Acad Sci USA 2017, 114, 11968.

(26) Li, X.; Gu, C.; Hostachy, S.; Sahu, S.; Wittwer, C.; Jessen, H. J.; Fiedler, D.; Wang, H.; Shears Stephen, B. Control of XPR1-dependent cellular phosphate efflux by InsP8 is an exemplar for functionally-exclusive inositol pyrophosphate signaling. Proc Natl Acad Sci USA 2020, 117, 3568.

(27) Wilson, M. S.; Jessen, H. J.; Saiardi, A. The inositol hexakisphosphate kinases IP6K1 and -2 regulate human cellular phosphate homeostasis, including XPR1-mediated phosphate export. J Biol Chem 2019, 294, 11597.

(28) Moritoh, Y.; Abe, S.-i.; Akiyama, H.; Kobayashi, A.; Koyama, R.; Hara, R.; Kasai, S.; Watanabe, M. The enzymatic activity of inositol hexakisphosphate kinase controls circulating phosphate in mammals. Nat Commun 2021, 12, 4847.

(29) Yousaf, R.; Gu, C.; Ahmed, Z. M.; Khan, S. N.; Friedman, T. B.; Riazuddin, S.; Shears, S. B.; Riazuddin, S. Mutations in Diphosphoinositol-Pentakisphosphate Kinase PPIP5K2 are associated with hearing loss in human and mouse. PLoS genetics 2018, 14, e1007297.

(30) Khaled, M. L.; Bykhovskaya, Y.; Gu, C.; Liu, A.; Drewry, M. D.; Chen, Z.; Mysona, B. A.; Parker, E.; McNabb, R. P.; Yu, H.; Lu, X.; Wang, J.; Li, X.; Al-Muammar, A.; Rotter, J. I.; Porter, L. F.; Estes, A.; Watsky, M. A.; Smith, S. B.; Xu, H.; Abu-Amero, K. K.; Kuo, A.; Shears, S. B.; Rabinowitz, Y. S.; Liu, Y. PPIP5K2 and PCSK1 are Candidate Genetic Contributors to Familial Keratoconus. Sci Rep 2019, 9, 19406.

(31) Du, C.; Wang, X.; Wu, Y.; Liao, W.; Xiong, J.; Zhu, Y.; Liu, C.; Han, W.; Wang, Y.; Han, S.; Chen, S.; Xu, Y.; Wang, S.; Wang, F.; Yang, K.; Zhao, J.; Wang, J. Renal Klotho and inorganic phosphate are extrinsic factors that antagonistically regulate hematopoietic stem cell maintenance. Cell Rep 2022, 38, 110392.

(32) Stephens, L.; Radenberg, T.; Thiel, U.; Vogel, G.; Khoo, K. H.; Dell, A.; Jackson, T. R.; Hawkins, P. T.; Mayr, G. W. The detection, purification, structural characterization, and metabolism of diphosphoinositol pentakisphosphate(s) and bisdiphosphoinositol tetrakisphosphate(s). J Biol Chem 1993, 268, 4009.

(33) Lin, H.; Fridy, P. C.; Ribeiro, A. A.; Choi, J. H.; Barma, D. K.; Vogel, G.; Falck, J. R.; Shears, S. B.; York, J. D.; Mayr, G. W. Structural Analysis and Detection of Biological Inositol Pyrophosphates Reveal That the Family of VIP/Diphosphoinositol Pentakisphosphate Kinases Are 1/3-Kinases*. J Biol Chem 2009, 284, 1863.

(34) Blüher, D.; Laha, D.; Thieme, S.; Hofer, A.; Eschen-Lippold, L.; Masch, A.; Balcke, G.; Pavlovic, I.; Nagel, O.; Schonsky, A.; Hinkelmann, R.; Wörner, J.; Parvin, N.; Greiner, R.; Weber, S.; Tissier, A.; Schutkowski, M.; Lee, J.; Jessen, H.; Schaaf, G.; Bonas, U. A 1-phytase type III effector interferes with plant hormone signaling. Nat Commun 2017, 8, 2159.

(35) Ito, M.; Fujii, N.; Wittwer, C.; Sasaki, A.; Tanaka, M.; Bittner, T.; Jessen, H. J.; Saiardi, A.; Takizawa, S.; Nagata, E. Hydrophilic interaction liquid chromatography–tandem mass spectrometry for the quantitative analysis of mammalian-derived inositol poly/pyrophosphates. J Chromatogr A 2018, 1573, 87.

(36) Kobayashi, A.; Abe, S.-i.; Watanabe, M.; Moritoh, Y. Liquid chromatography-mass spectrometry measurements of blood diphosphoinositol pentakisphosphate levels. J Chromatogr A 2022, 1681, 463450.

(37) Harmel, R. K.; Puschmann, R.; Nguyen Trung, M.; Saiardi, A.; Schmieder, P.; Fiedler, D. Harnessing 13C-labeled myo-inositol to interrogate inositol phosphate messengers by NMR. Chem Sci 2019, 10, 5267.

(38) Qiu, D.; Wilson, M. S.; Eisenbeis, V. B.; Harmel, R. K.; Riemer, E.; Haas, T. M.; Wittwer, C.; Jork, N.; Gu, C.; Shears, S. B.; Schaaf, G.; Kammerer, B.; Fiedler, D.; Saiardi, A.; Jessen, H. J. Analysis of inositol phosphate metabolism by capillary electrophoresis electrospray ionization mass spectrometry. Nat Commun 2020, 11, 6035.

(39) Haas, T. M.; Mundinger, S.; Qiu, D.; Jork, N.; Ritter, K.; Durr-Mayer, T.; Ripp, A.; Saiardi, A.; Schaaf, G.; Jessen, H. J. Stable Isotope Phosphate Labelling of Diverse Metabolites is Enabled by a Family of (18) O-Phosphoramidites. Angew Chem Int Ed 2022, 61, e202112457.

(40) Qiu, D.; Eisenbeis, V. B.; Saiardi, A.; Jessen, H. J. Absolute Quantitation of Inositol Pyrophosphates by Capillary Electrophoresis Electrospray Ionization Mass Spectrometry. JoVE 2021, e62847.

(41) Gu, C.; Wilson, M. S. C.; Jessen, H. J.; Saiardi, A.; Shears, S. B. Inositol Pyrophosphate Profiling of Two HCT116 Cell Lines Uncovers Variation in InsP8 Levels. Plos One 2016, 11, e0165286.

(42) Draškovic, P.; Saiardi, A.; Bhandari, R.; Burton, A.; Ilc, G.; Kovacevic, M.; Snyder, S. H.; Podobnik, M. Inositol Hexakisphosphate Kinase Products Contain Diphosphate and Triphosphate Groups. Chem Biol 2008, 15, 274.

(43) Puschmann, R.; Harmel, R. K.; Fiedler, D. Scalable Chemoenzymatic Synthesis of Inositol Pyrophosphates. Biochemistry-Us 2019, 58, 3927.

(44) Capolicchio, S.; Thakor, D. T.; Linden, A.; Jessen, H. J. Synthesis of unsymmetric diphospho-inositol polyphosphates. Angew Chem Int Ed 2013, 52, 6912.

(45) Capolicchio, S.; Wang, H.; Thakor, D. T.; Shears, S. B.; Jessen, H. J. Synthesis of densely phosphorylated bis-1,5-diphospho-myo-inositol tetrakisphosphate and its enantiomer by bidirectional P-anhydride formation. Angew Chem Int Ed 2014, 53, 9508.

(46) Bittner, T.; Wittwer, C.; Hauke, S.; Wohlwend, D.; Mundinger, S.; Dutta, A. K.; Bezold, D.; Dürr, T.; Friedrich, T.; Schultz, C.; Jessen, H. J. Photolysis of Caged Inositol Pyrophosphate InsP8 Directly Modulates Intracellular Ca2+ Oscillations and Controls C2AB Domain Localization. J Am Chem Soc 2020, 142, 10606.

(47) Hauke, S.; Dutta, A. K.; Eisenbeis, V. B.; Bezold, D.; Bittner, T.; Wittwer, C.; Thakor, D.; Pavlovic, I.; Schultz, C.; Jessen, H. J. Photolysis of cell-permeant caged inositol pyrophosphates controls oscillations of cytosolic calcium in a β-cell line. Chem Sci 2019, 10, 2687.

