## supplementary file for "Capillary electrophoresis mass spectrometry identifies new isomers of inositol pyrophosphates in mammalian tissues"

### Supplementary Methods

#### Supplementary method 1: PPIP5K2 knockout mouse generation

A single CAS9 target site (SgA: ACCGTTTGCTATTTAAAACAAGG[PAM]) was utilized to generate a double stranded break in the *Ppip5k2* locus at the splice acceptor of exon 9. PPIP5K2-SgA SgRNA was ordered from IDTDNA (Coralville, IA, USA). EnGen Spy Cas9 NLS nuclease was purchased from New England Biolabs (Ipswich. Massachusetts, USA). C57BL/6J one-cell embryos were microinjected with CAS9 protein (70 ng/ul) and SgRNA-A (40 ng/ul). Microinjected embryos were surgically transferred to SWISS pseudo-pregnant females. At weaning, potential founders were screened by PCR amplicon sequencing (FWD: 5'-TCAAATGGAAGTAATGCAACCTGC-3'; Rev 5'-TCAAAGTCTTCTCTGTGGCTCAA-3'). Multiple founders were identified with NHEJ mutations that likely disrupt PPIP5K2 expression. Founders of interest were bred to wildtype C57BL/6J mice and F1 offspring were re-screened to confirm germline transmission of the mutant allele. A founder line with a 21 bp NHEJ deletion (ttgttttaaatag-CAAACGGT) at the intron-exon (lowercase-uppercase) boundary was selected as the most likely line to genetically disrupt *Ppip5k2*. The mutant mouse line was crossed to wildtype C57BL/6J mice for at least two generations to eliminate any unknown, unlinked mutations. Computationally, no likely genetically linked mutations were predicted near the target site. The PPIP5k2-deficient mouse colony was genotyped by Transnetyx using primer/probe assays (Fwd: TGGATTCATAGTAGTTCCAATGATACCCTAA, Rev: ATAGGATTGTCCATTGGCACGTAA, WT-Probe: ACAAACCGTTTGCTATTTAA, Null-Probe: CAAATCAAAGCCACAAGGTA).

Once mice homozygous for the d(21) allele were generated, splicing from exon 8 to exon 10 was confirmed by cDNA amplicon sequencing (cDNA-Fwd: ACCGTGATCCAAATAACCCGA; cDNA-Rev: TTTTCGGACAGCCATGAGGAA). The predominant splice variant identified has exon 8 to exon 10 splicing, which results in a nonsense mutation followed by premature stop in 22 codons. A minor mutant splice variant was identified with exon 8 splicing to the middle of exon 9, which results in a nonsense mutation with a premature stop in 10 codons. Neither mutant transcript variant is expected to be yield a functional protein.

#### Supplementary method 2: PBMCs

Lymphocyte isolation

Healthy donor PBMC samples were recruited from the HBUF biobank of the University Medical Center Freiburg with approval of the Institutional Review Boards (Ethics committee of the Albert-Ludwigs-University, Freiburg, #407/16).

Peripheral blood mononuclear cells (PBMCs) were isolated from whole blood samples using density gradient centrifugation. Whole blood was diluted 1:1 with 1× phosphate-buffered saline (PAN Biotech GmbH (Aidenbach, Germany)), layered on top of Pancoll separation medium (PAN Biotech GmbH (Aidenbach, Germany), and) centrifuged at 1200*g* for 20 min. The PBMC layer was retrieved, resuspended in 20 ml of PBS, and centrifuged again at 300*g* for 10 min. Cells were then resuspended in fetal calf serum with 10% DMSO (fetal calf serum, Thermo Scientific (Waltham, MA); DMSO, PanReac AppliChem) and stored at -80°C until used.

PBMCs were thawed in RPMI 1640 medium supplemented with 10% fetal calf serum, 1% penicillin/streptomycin, and 1.5% HEPES buffer 1 mol/L (complete medium; all additives from Thermo Scientific (Waltham, MA)) and incubated for 15–30 min at 37 °C in complete medium containing 50 U $ml^{-1}$ nuclease (Thermo Scientific (Waltham, MA)) before processing.

For CD8+ T cell isolation $1x10^9$ PBMCs were resuspended in PBS supplemented with 2% fetal calf serum and 0,1% EDTA buffer 0,5 M (PBS, PAN Biotech GmbH (Aidenbach, Germany)); all additives from Thermo Scientific (Waltham, MA)). Cells were labeled by adding antibody complexes recognizing CD8 and magnetic particles (EasySep Human CD8 positive Selection Kit II (STEMCELL)). Labeled cells were isolated by using an EasySep magnet with a purity of 70-90%.

**Supplementary method 3: Colon biopsy from patient**

Study participants undergoing diagnostic screening colonoscopy donating colorectal biopsy samples of macroscopically normal mucosa at the endoscopy unit of the University Hospital of Freiburg were recruited from the HBUF biobank of the University Medical Center Freiburg (Ethics committee of the Albert-Ludwigs-University, Freiburg, #407/16 & #14/17). Patients undergoing diagnostic screening colonoscopy were recruited following patients' informed consent, intestinal biopsies were collected and tissue samples were frozen at -80°C until used.

**Supplementary method 4: Extraction of PP-InsPs and InsPs from mouse tissues and mouse feces**

Mouse tissue and feces were collected and frozen in liquid nitrogen immediately after mice were CO2 euthanized. For liver samples, the largest (i.e., the left) lobe was sampled. One lung (from the left side) was always sampled. Remaining procedures were performed at 4°C. Tissues were weighed and homogenized in 1 ml of 1M perchloric acid. The homogenate was centrifuged (13,500 x g; 10 min), then the supernatant was incubated with 10mg TiO2 beads for 30min to capture inositol pyrophosphates. The beads were washed with water and then inositol phosphates were eluted with 10% ammonium hydroxide by incubation for 20 min. The elute were dried in a vacuum centrifuge and stored at -80°C.

**Supplementary method 5: Extraction of PP-InsPs and InsPs from mouse food, PBMCs and colon biopsy**.

**Mouse food:** mouse food was powdered with a grinder. 40 mg of mouse food powder was then resuspended in 1 ml of ice-cold 1M perchloric acid solution and incubated at 4°C with rotation for 20 minutes. The suspension was centrifuged and the supernatant was transferred to pre-washed titanium dioxide ($TiO_2$) beads for purification according to the literature (Bio Protoc. 2018, 8, e2959). Briefly, the supernatant was incubated with the $TiO_2$ beads at 4°C with rotation for 20 minutes to allow adequate binding of the PP-InsPs and InsPs to the beads. After washing with 0.5 ml of 1M perchloric acid solution twice, 3% ammonium hydroxide was used to elute PP-InsPs and InsPs. The elution concentrated with a speed vac evaporator at 60°C for 1-3 h.

**PBMCs:** 12 million CD8+ T cells or 80 million depleted cells were collected and washed with PBS buffer, resuspended in 1 ml of ice-cold 1M perchloric acid solution. After incubating at 4°C with rotation for 20 minutes, the cell suspension was centrifuged and the supernatant was transferred to $TiO_2$ purification as described in the extraction of mouse food.

**Colon biopsy:** thaw the colon biopsy sample on ice and rinse it in ice-cold PBS. Quickly dry the colon biopsy with paper tissues to remove the excessive PBS, and then weigh the biopsy sample. The biopsy was homogenized with glass pestle tissue grinders in 1M perchloric acid solution. Keep the biopsy on ice for the whole procedure. After homogenizing, transfer the sample to 4°C with rotation for 20 minutes. Then the suspension was centrifuged and the supernatant was transferred to TiO2 purification as described in the extraction of mouse food.

**Supplementary method 6: Use of stable isotopic internal standards**

Quantitation of PP-InsP and InsP in this study was performed with known amounts of individual isotopic standards (IS) spiked into the samples. Usually, ratio of analyte peak area $(Area)^{12C}$ / IS peak area $(Area)^{13C}$ is <5 to ensure a linear relationship to analyte concentration/IS concentration. Due to the wide range of PP-InsP and InsP levels across mouse tissues, we obtained ratios of analyte peak area $(Area)^{12C}$ / IS peak area $(Area)^{13C}$ in the proper range, simply by giving tissue samples different dilution factors. An internal standard mixture was prepared, containing 2 µM [$^{13}C_6$] 1, 5-$(PP)_2$-$InsP_5$, 8 µM [$^{13}C_6$] 5-PP-$InsP_5$, 4 µM [$^{13}C_6$] 1-PP-$InsP_5$, 40 µM [$^{13}C_6$]$InsP_6$ and 40 µM [$^{13}C_6$]2-OH $InsP_5$. All mouse tissue extracts were dissolved in 30µL water, also applied for mouse feces and food extracts. For heart, kidney, lung and spleen, 7.5 µL IS mixture and 7.5 µL sample solution were mixed in CE vial. For liver and colon, 7.5 µL IS mixture, 3 µL sample solution and 4.5 µL water were mixed. For mouse feces and food, 10 µL IS mixture, 1 µL sample solution and 9 µL water were mixed.

**Supplementary method 7: CE-QQQ analysis**

A CE-ESI-QQQ system is used in this study, which consists of an Agilent 7100 CE, a triple quadrupole tandem mass spectrometry Agilent 6495c, hyphenated by an Agilent Jet Stream (AJS) electrospray ionization (ESI) source. Commercial CE-MS sheath liquid coaxial interface was adopted, with an isocratic LC pump constantly delivering the sheath-liquid (via a splitter with a ratio of 1:100).

All experiments were performed on a bare fused silica capillary with a length of 100 cm (50 µm internal diameter and 365 µm outer diameter). Two background eletrolytes (BGE) were employed for this study. BGE A is 35mM ammonium acetate titrated by ammonia solution to pH 9.7. BGE B is 40mM ammonium acetate titrated by ammonia solution to pH 9.0. Capillary was flushed by BGE for 400s between runs. Samples were injected by applying 100 mbar pressure for 10 s, corresponding to 1 % of the total capillary volume (20 nL).

The sheath liquid is a mixture of water-isopropanol (1/1, v/v) and with a constant flow at 10 µL/min. The MS source parameters settings: nebulizer pressure was 8 psi, gas temperature was 150 °C and with a flow of 11 L/min, sheath gas temperature was 175 °C and with a flow at 8 L/min. Capillary voltage was -2000 V with nozzle voltage 2000V. Negative high-pressure RF and low-pressure RF (Ion Funnel parameters) was 70V and 40V, respectively. Mass spectrometer parameters for MRM transitions are shown below.

| **Compound Name** | **Precursor Ion** | **Product Ion** | **dwell** | **Frag (V)** | **CE (V)** | **Cell Acc (V)** | **Polarity** |
|---|---|---|---|---|---|---|---|
| $InsP_8$ | 408.9 | 359.9 | 50 | 166 | 9 | 1 | Negative |
| $[^{13}C_6]InsP_8$ | 411.9 | 362.9 | 50 | 166 | 9 | 1 | Negative |
| $InsP_7$ | 368.9 | 319.9 | 50 | 166 | 9 | 3 | Negative |
| $[^{13}C_6]InsP_7$ | 371.9 | 322.9 | 50 | 166 | 9 | 3 | Negative |
| $[^{18}O_2]InsP_7$ | 370.9 | 319.9 | 50 | 166 | 9 | 3 | Negative |
| $InsP_6$ | 328.9 | 480.9 | 50 | 166 | 13 | 4 | Negative |
| $[^{13}C_6]InsP_6$ | 331.9 | 486.9 | 50 | 166 | 13 | 4 | Negative |
| $InsP_6$ | 328.9 | 79.1 | 50 | 166 | 53 | 4 | Negative |
| $[^{13}C_6]InsP_6$ | 331.9 | 79.1 | 50 | 166 | 53 | 4 | Negative |
| $InsP_5$ | 289 | 498.9 | 50 | 166 | 9 | 3 | Negative |
| $[^{13}C_6]InsP_5$ | 292 | 504.9 | 50 | 166 | 9 | 3 | Negative |

### *Supplementary Figures*

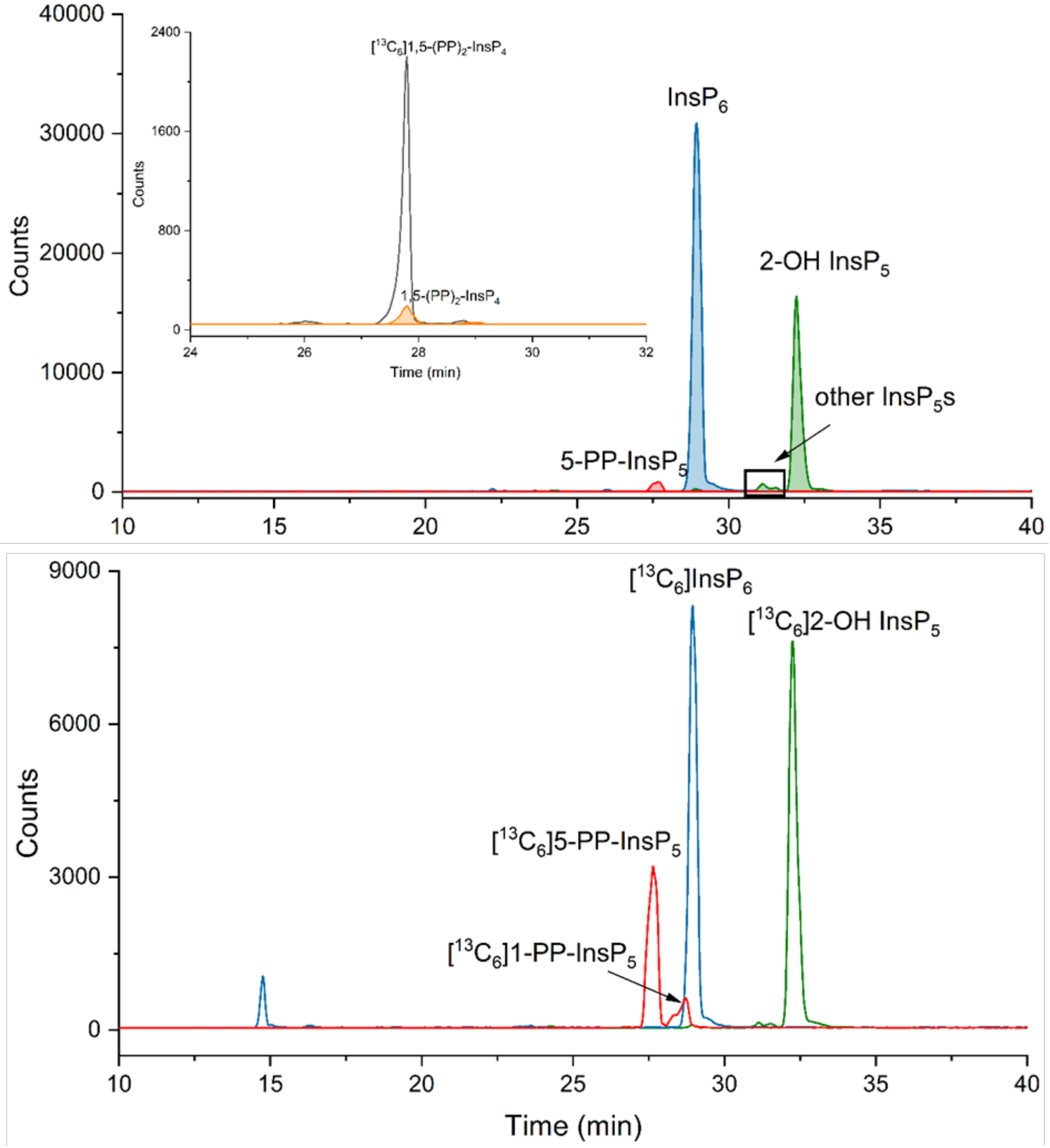


**Supplementary Figure 1** The upper panel shows extracted ion electropherograms (EIEs) of PP-InsPs and InsPs prepared from mouse liver by CE-QQQ using BGE-A. The upper panel shows the migration of 5-PP-$InsP_5$ (red), $InsP_6$ (blue) and $InsP_5$ (green) with filled plots. The inset depicts the co-elution of 1,5-$(PP)_2$-$InsP_4$ (filled orange plot) spiked with an internal standard of 1 μM [$^{13}C_6$] 1,5-$(PP)_2$-$InsP_4$ (black plot). The lower panel shows the migration of 4 μM [$^{13}C_6$] 5-PP-$InsP_5$, 2 μM [$^{13}C_6$] 1-PP-$InsP_5$, 20 μM [$^{13}C_6$]$InsP_6$ and 20 μM [$^{13}C_6$]2-OH $InsP_5$ spiked into the sample solution of PP-InsPs and InsPs. The PP-InsP and InsP species were assigned by multiple reaction monitoring (MRM) and identical migration time with spiked stable isotope labeled (SIL) standards. Accurate masses of 5-PP-$InsP_5$, $InsP_6$, 2-OH $InsP_5$ and other $InsP_5$s were also confirmed by CE-qTOF analysis.

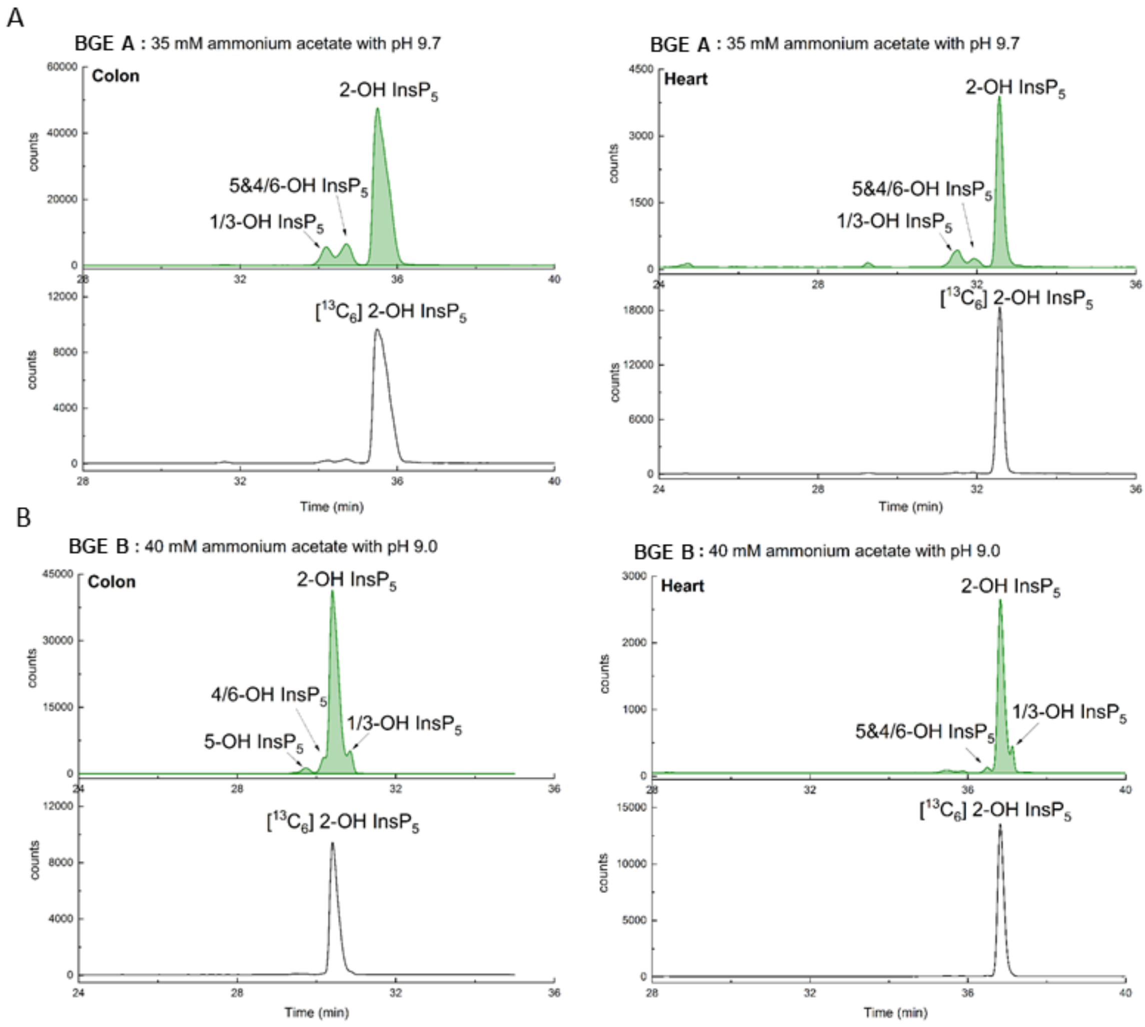


**Supplementary Figure 2** $InsP_5$ isomer identification with [$^{13}C_5$]2-OH $InsP_5$ (black line) in mouse colon and heart by CE-QQQ. **(a)** EIEs of $InsP_5$ using BGE A. Same as our earlier study, 1/3-OH $InsP_5$ migrated firstly, then 5&4/6-OH $InsP_5$, and 2-OH $InsP_5$ migrated in the end (green trace). 2-OH $InsP_5$ is main $InsP_5$ isomer in mouse tissue. **(b)** EIEs of $InsP_5$ using BGE B. The migration order of $InsP_5$ isomers is 5-OH $InsP_5$, 4/6-OH $InsP_5$, 2-OH $InsP_5$ and 1/3-OH $InsP_5$ (green trace).

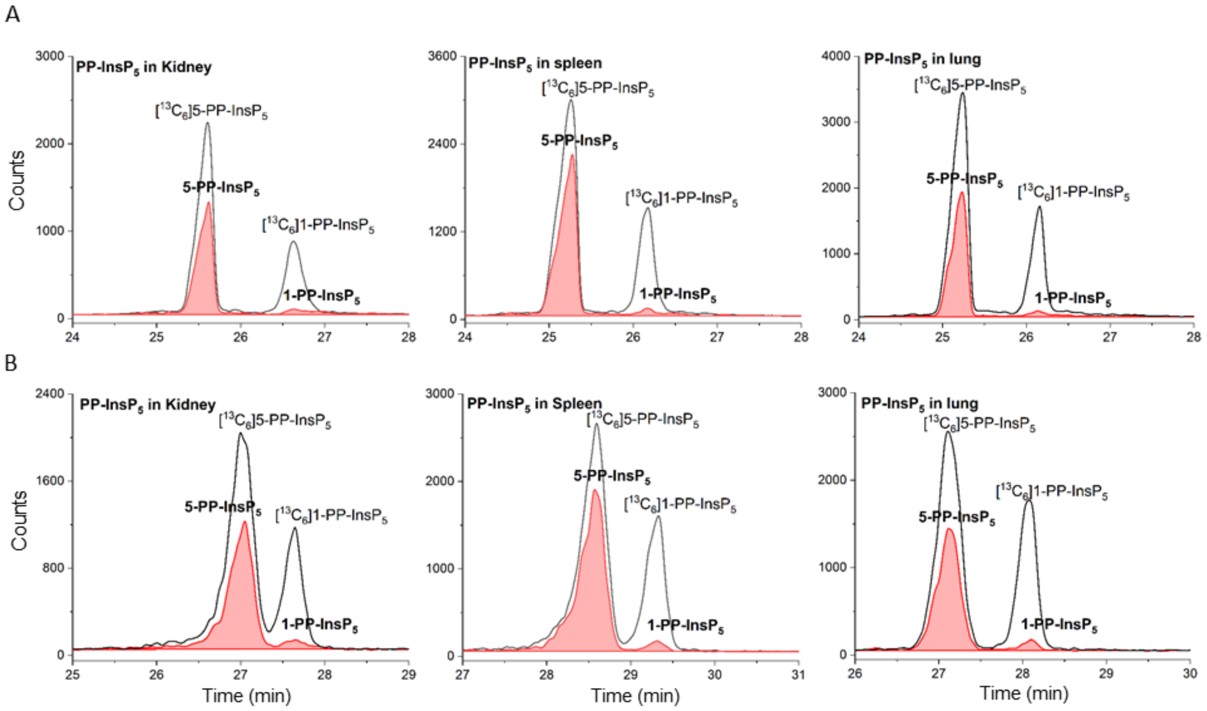


**Supplementary Figure 3** Canonical 5-PP-InsP$_5$ and 1-PP-InsP$_5$ were found in mouse kidney, spleen and lung, same as in mouse liver. **(A)** Extracted ion electropherograms (EIEs) of PP-InsP$_5$ (red trace) in mouse kidney, spleen and lung with stable isotope ($^{13}$C) labeled internal reference compounds (black line) using BGE A containing 35 mM ammonium acetate titrated with ammonium hydroxide to pH 9.7 as in Figure 2C. 2-PP-InsP$_5$ migrated after 1-PP-InsP$_5$ and would be identified under this condition. **(B)** Extracted ion electropherograms (EIEs) of PP-InsP$_5$ in mouse kidney, spleen and lung (red trace) with stable isotope ($^{13}$C) labeled internal reference compounds (black line) using BGE B containing 40 mM ammonium acetate titrated with ammonium hydroxide to pH 9.0 as in Figure 2D. 4/6-PP-InsP$_5$ migrated closely after 5-PP-InsP$_5$ and would be identified under this condition.

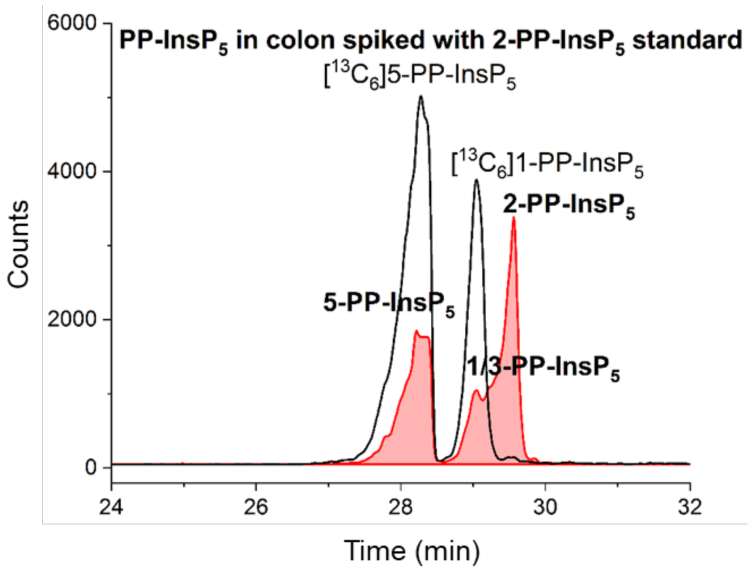


**Supplementary Figure 4** PP-Ins$P_5$ in mouse colon tissue spiked with 2-PP-Ins$P_5$ standard with BGE A containing 35 mM ammonium acetate titrated with ammonium hydroxide to pH 9.7. The PP-Ins$P_5$ isomer in colon migrated after 1/3-PP-Ins$P_5$ has same migration time as the 2-PP-Ins$P_5$ standard.

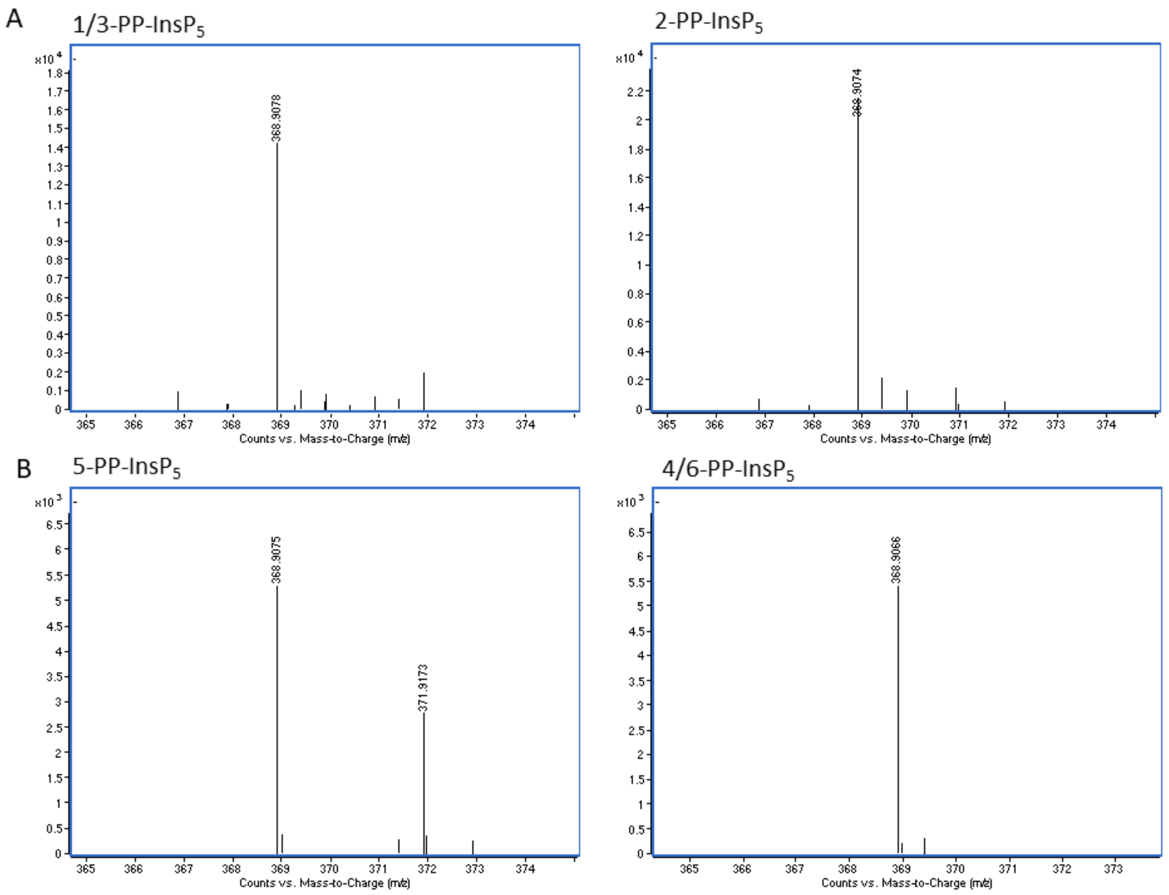


**Supplementary Figure 5** High resolution mass spectra (HRMS) of PP-Ins$P_5$ in mouse colon tissue. (**A**) HRMS of 1/3-PP-Ins$P_5$ and 2-PP-Ins$P_5$ in mouse colon tissue with [$^{13}C_6$]5-PP-Ins$P_5$ reference from CE-qTOF analysis using BGE A. Theoretical mass to charge (m/z, z=2) value for PP-Ins$P_5$ and [$^{13}C_6$]PP-Ins$P_5$ is 368.9066 and 371.9166, respectively. (**B**) HRMS of 5-PP-Ins$P_5$ and 4/6-PP-Ins$P_5$ in mouse colon tissue with [$^{13}C_6$]1-PP-Ins$P_5$ reference from CE-qTOF analysis using BGE B.

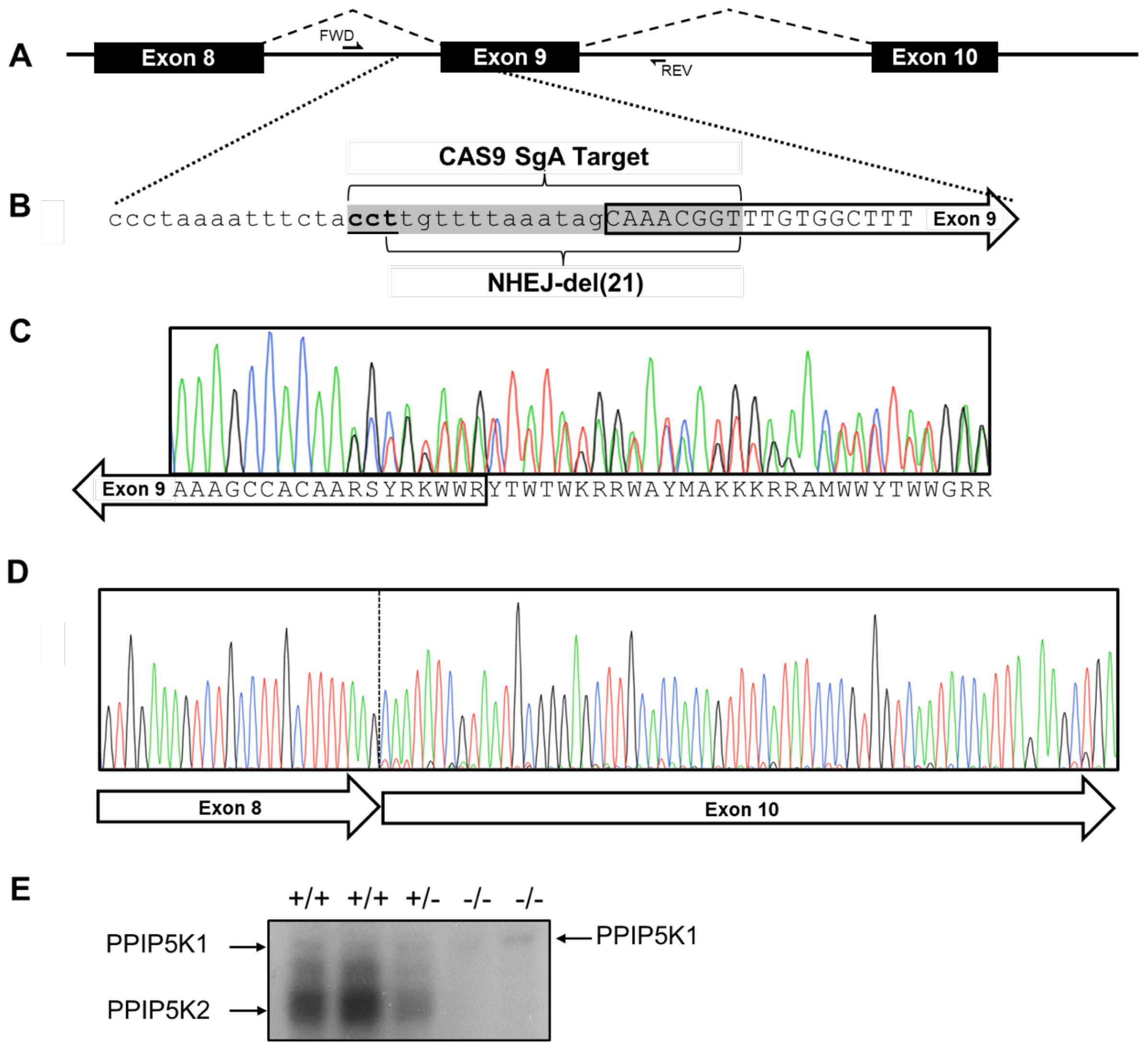


**Supplementary Figure 6** PPIP5K2 CRISPR/Cas9 Targeting Scheme. A) Endogenous Ppip5k2 locus region around exon 9. B) Nucleotide level resolution with annotation exon 9 (solid arrow), CAS9 target sequence (gray) with PAM (underlined). C) Genomic sequencing in F1 mice that are heterozygous for the 21 bp NHEJ deletion, which disrupts the canonical splicing from exon 8 to exon 9. D) Homozygous mutant cDNA transcript sequencing confirmed that the primary splicing event was exon 8 to exon 10, which results in a nonsense mutation and a premature stop in 22 codons. A secondary minor splice event was identified with exon 8 splicing to the middle of exon 9, which results in a nonsense mutation and a premature stop in 10 codons. Neither mutant splice event is expected to yield a functional protein. E) PPIP5K2 protein cannot be detected in the liver of PPIP5K2 KO mice. The faint PPIP5K1 band is due to cross reactivity with PPIP5K2 antibody

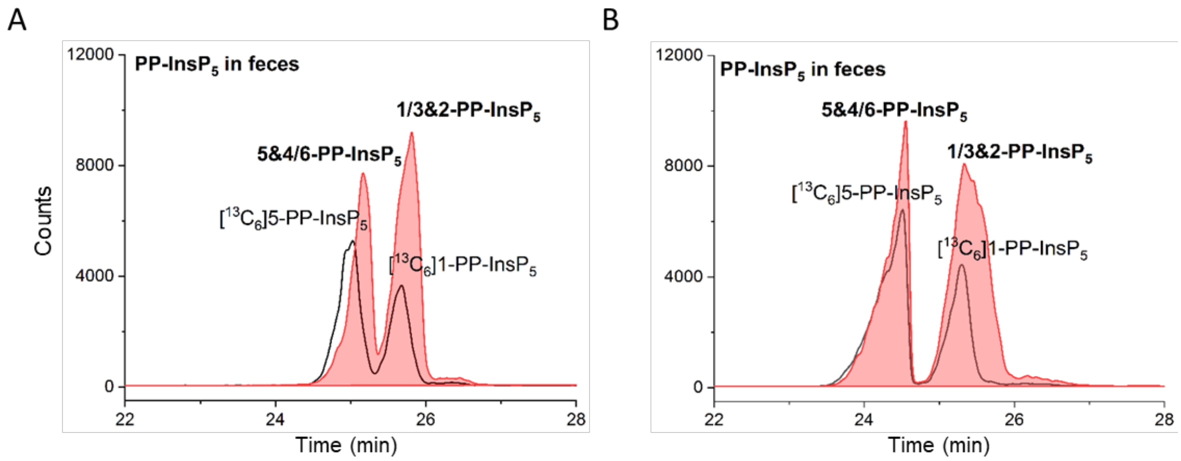


**Supplementary Figure 7** CE-QQQ analysis of PP-$InsP_5$ isomer distribution in mouse feces with stable isotope ($^{13}C$) labeled internal reference compounds (black line) **(A)** EIEs of PP-$InsP_5$ with BGE A, showing the existence of 4/6-PP-$InsP_5$ in mouse feces and its abundance is higher than 5-PP-$InsP_5$ **(B)** EIEs of PP-$InsP_5$ with BGE B. 2-PP-$InsP_5$ is apparently present in feces.

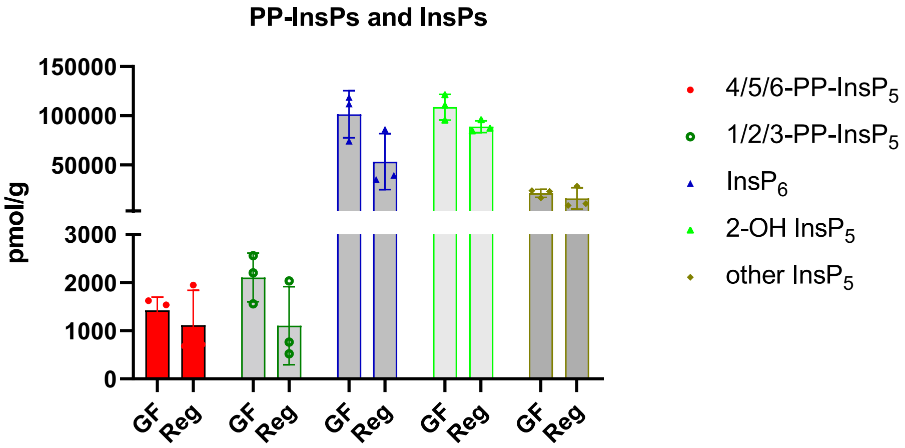


**Supplementary Figure 8** Comparison of PP-InsPs and InsPs in germ-free (GF) and regular (Reg) mouse colon samples.

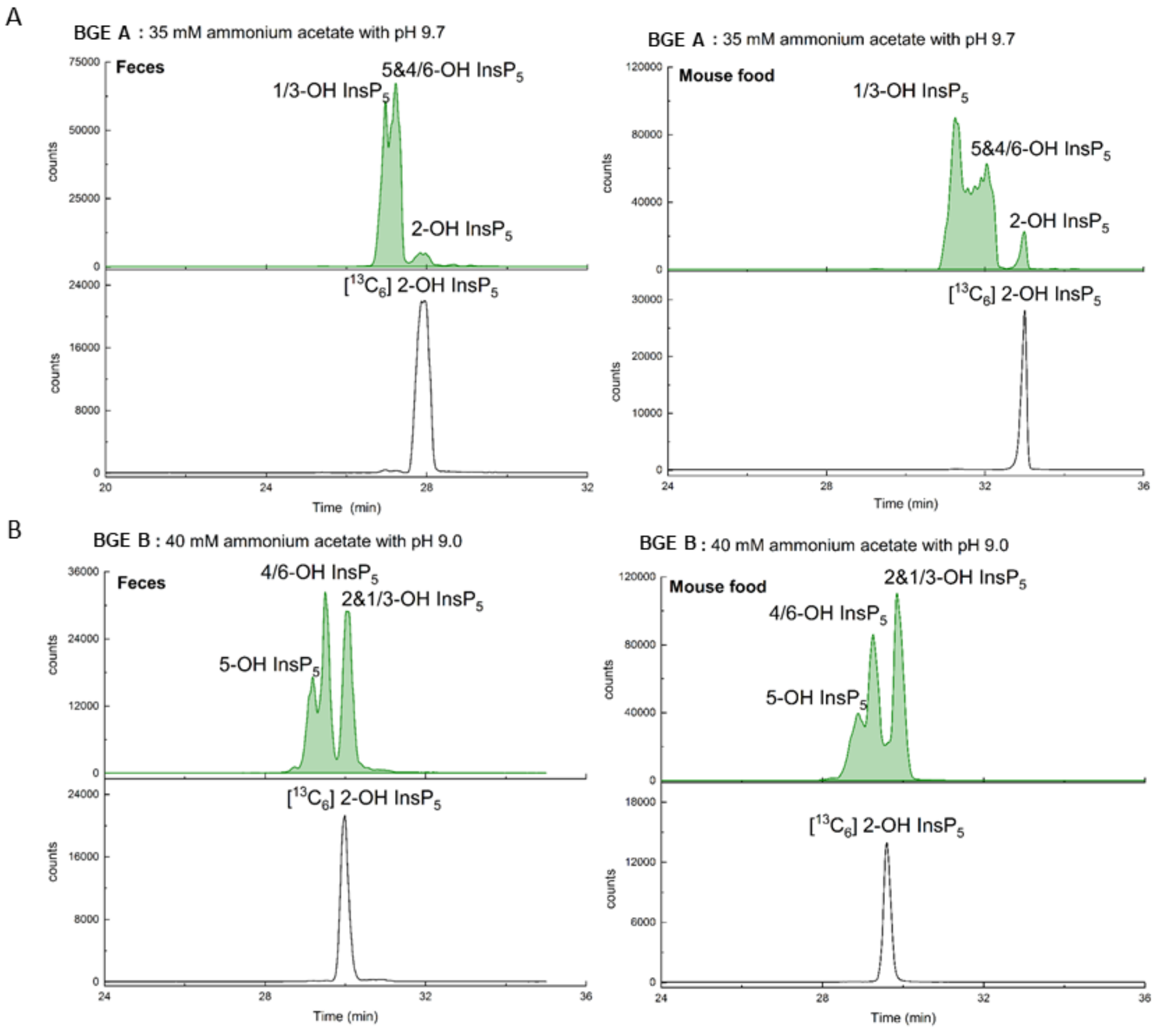


**Supplementary Figure 9** $InsP_5$ isomer distribution in mouse feces and mouse food by CE-QQQ. **(A)** EIEs of $InsP_5$ and [$^{13}C_5$]2-OH $InsP_5$ using BGE A, 2-OH $InsP_5$ is the minor $InsP_5$ isomer in both mouse feces and mouse food, while 2-OH $InsP_5$ is the major $InsP_5$ isomer in mouse tissues (Figure 2A). **(B)** EIEs of $InsP_5$ and [$^{13}C_5$]2-OH $InsP_5$ using BGE B, 5-OH $InsP_5$, 4/6-OH $InsP_5$ and 1/3-OH $InsP_5$ all substantially contribute to the $InsP_5$ pools in mouse feces and mouse food.

**Synthesis of $^{18}O$ labeled 4-PP-InsP$_5$.**

**Chemical Synthesis**

### 1. General Remarks

**Reactions** were carried out using flame-dried glassware under an atmosphere of dry Argon and magnetically stirred, unless noted otherwise. Air- and moisture-sensitive liquids and solutions were transferred via syringe or stainless steel canula.

**$H_2{}^{18}O$** (>99% purity, 99% isotopic enrichment) was purchased from Taiyo Nippon Sanso (respectively from Cortecnet as european vendor).

**Reagents** were purchased from commercial suppliers (Acros, Aldrich, Fluka, TCI) and used without further purification, unless noted otherwise.

**Solvents** were obtained in analytical grade and used as received for extractions, precipitation and solid washing.

**Dry solvents** for reactions were purchased in a dry form from Sigma and stored over molecular sieves as well as under the atmosphere of dry $N_2$.

**Deuterated solvents** for NMR and reactions were obtained from Armar Chemicals, Switzerland and Eurisotop, Germany, in the indicated purity grade and used as received for NMR spectroscopy.

**Strong ion-exchange chromatography** was performed using an automated Äkta® – system. Q-Sepharose was purchased from Aldrich. Buffer solutions were produced manually using milliQ $H_2O$.

**Preparative RP-MPLC** was performed using an automated Interchim® - system. The AQ-solid phase was purchased from Interchim.

**Lyophilizations** were done with Christ Freeze Dryer Alpha 1-4 LDplus and Christ Freeze Dryer Alpha 1-2 LDplus.

**Analytical HPLC-MS** measurements were performed using a Thermofisher Ultimate 3000 system coupled to MSQplus. C18-AQ-columns were purchased from ProntoSil.

**$^{1}$H-NMR spectra** were recorded on Bruker 300 MHz spectrometers and Bruker 400 MHz (with cryoprobe) spectrometer in the indicated deuterated solvent. Data are reported as follows: chemical shift (δ, ppm), multiplicity (s, singlet; d, doublet; t, triplet; q, quartet; m, multiplet; br. s, broad signal), coupling constant(s) (J, Hz), integration. All signals were referenced to the internal solvent signal as standard ($D_2O$, δ 4.79; MeCN-$d_3$, δ 1.94, $CDCl_3$, 7.26).

**$^{13}$C{$^{1}$H}-NMR spectra** were recorded with $^{1}$H-decoupling on Bruker 126 MHz, Bruker 101 MHz (with cryoprobe) spectrometers at 298K in the indicated deuterated solvent.

**$^{31}$P{$^{1}$H}-NMR spectra and $^{31}$P-NMR spectra** were recorded with $^{1}$H-decoupling or $^{1}$H coupling, respectively, on Bruker 162 MHz (with cryoprobe) and Bruker 122 MHz spectrometers in the indicated deuterated solvent. All signals were referenced to an internal standard (PPP).

**Mass spectra** were recorded by C. Warth (Mass spectrometry service of the University of Freiburg) on a Thermo LCQ Advantage [spray voltage: 2.5 – 4.0 kV, spray current: 5 µA, ion transfer tube: 250 (150) °C, evaporation temperature: 50 – 400°C.

**Synthesis of AB-Phosphoramidite S1**

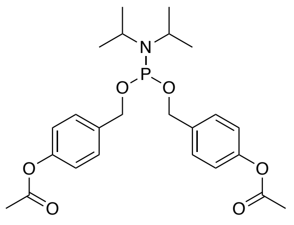


**S1**

$C_{24}H_{32}NO_6P$
461,49

The compound was synthesized according to Jessen et al.[1] Analytical data are in accordance with literature.

#### Synthesis of Inositol Monophosphate S2

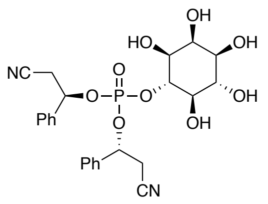


**S2**

$C_{24}H_{27}N_2O_9P$
MW: 518,46

The compound was synthesized according to Capolicchio et al.[2] Analytical data are in accordance with literature.

#### Synthesis of $^{18}O_2$-AB-PMB-P-amidite S3

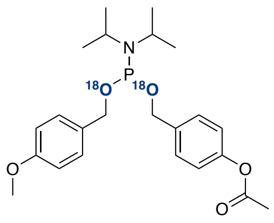


**S3**

$C_{23}H_{32}NO_3{}^{18}O_2P$
437,49

The compound was synthesized according to Haas et al.[3] Analytical data are in accordance with literature.

#### Synthesis of Hexakisphosphate 12

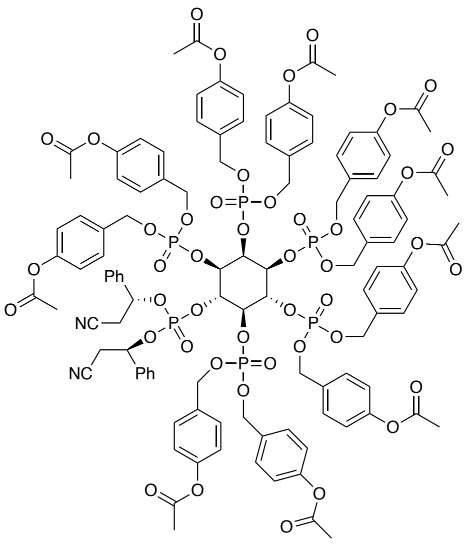

**12**

$C_{114}H_{112}N_2O_{44}P_6$
MW: 2399,96

Inositol monophosphate **S2** (350 mg, 675 µmol, 1.0 eq) and acyloxybenzyl phosphoramidite **S1** (3.12 g, 6.76 mmol, 10.0 eq) were coevaporated with MeCN (2 × 2 mL) and then dissolved in DMF (12 mL). 5-(Ethylthio)-1*H*-tetrazole (873 mg, 6.76 mmol, 10.0 eq) was added and the resulting mixture was stirred at r.t. The reaction progress was followed by $^{31}$P-NMR. After completion of the reaction (90 min), the mixture was cooled to 0°C and oxidation was achieved by addition of *m*CPBA (70% moistened with water, 1.67 g, 6.76 mmol, 10.0 eq) and stirring for 15 min at r.t. The mixture was diluted with EtOAc (300 mL) and washed with $H_2O$ (3 × 250 mL) and brine (250 mL). The solution was dried over $MgSO_4$ and the solvent was removed under reduced pressure. The residue was purified by an automated MPLC (Interchim-system C18-column, MeCN/$H_2O$) to obtain the title compound **12** (960 mg, 400 µmol, 59%) as a colorless oil.

**$^1$H-NMR** (400 MHz, $CDCl_3$ ): δ = 7.42 − 7.27 (m, 10H), 7.26 − 7.10 (m, 20H), 7.07 − 6.87 (m, 20H), 5.90 (td, *J* = 7.9, 4.3 Hz, 1H), 5.63 (td, *J* = 8.9, 4.0 Hz, 1H), 5.48 ($m_c$ , 1H), 5.30 – 4.89 (m, 22H), 4.29 (2H, $m_c$), 3.46 ($m_c$, 1H), 2.93 (dd, *J* = 16.7, 4.5 Hz, 1H), 2.82 – 2.67 (m, 2H), 2.61 − 2.50 (m, 1H), 2.31 – 2.19 (m, 27H), 2.09 (s, 3H) ppm. **$^{31}$P{$^1$H}-NMR** (162 MHz, $CDCl_3$): δ = -0.76 (s, 1P), -0.90 (s, 1P), -1.87 (s, 1P), -2.13 (s, 1P), -3.23 (s, 1P), -3.36 (s, 1P) ppm. **HRMS** (ESI) m/z for $C_{113}{}^{13}C_1H_{112}N_2NaP_6$ $[M+Na]^+$: calcd. 2422.4939 found 2422.4954.

### Synthesis of $^{18}$O-labeled Diphosphoinositol Pentakisphosphate 13

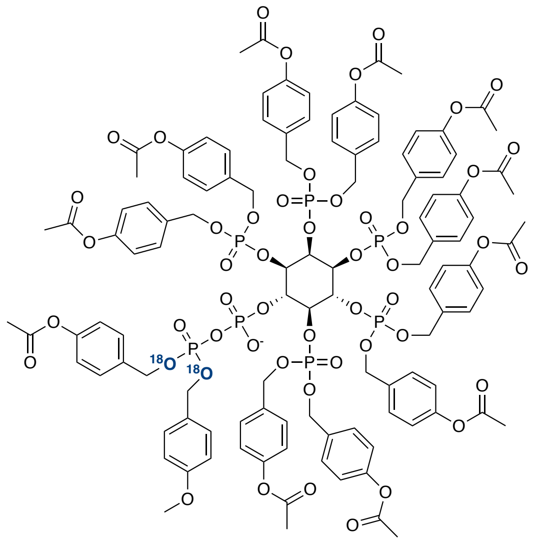

**13**

$C_{113}H_{114}O_{48}{}^{18}O_{2}P_{7}^{-}$
MW: 2492,92

Hexakisphosphate **12** (50.0 mg, 20.8 µmol, 1.0 eq.) was co-evaporated with MeCN (2 × 1.0 mL) and then dissolved in MeCN (2.0 mL). DBU (12.4 µL, 12.7 mg, 83.3 µmol, 4.0 eq.) and BSTFA (22.3 µL, 21.5 mg, 83.3 µmol, 4.0 eq.) were added. Turnover was monitored by TLC (EtOAc:MeOH, 9:1). After 8 minutes TFA (6.42 µL, 9.50 mg, 83.3 µmol, 4.0 eq.) in MeOH (20 µL) was added and the solvent was immediately removed under reduced pressure. The residue was resolved in MeCN (2 mL) and $^{18}$O-AB-PMB-P-amidite **S3** (18.2 mg, 41.7 µmol, 2.0 eq.) was added. A solution of ETT in MeCN (1m, 41.7 µL, 41.7 µmol, 2.0 eq.) was added and the reaction progress was followed by $^{31}$P-NMR. After completion of the reaction (30 min), *m*CPBA (70%, 10.3 mg, 41.7 µmol, 2.0 eq.) was added at 0°C and the mixture was stirred for 10 min at rt. The product was precipitated with $Et_2O$ (40 mL), redissolved in $CH_2Cl_2$ (2 mL) and reprecipitated twice with $Et_2O$ (45 mL). The precipitate was dried under high vacuum. The title compound was obtained as colorless solid and directly used without further purification and analysis.

### Synthesis of $^{18}$O-labeled Diphosphoinositol Pentakisphosphate 13a

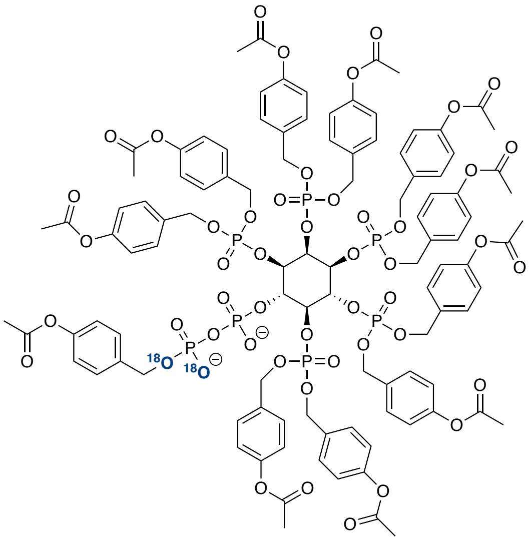


**13a**

$C_{105}H_{105}O_{47}{}^{18}O_2P_7{}^{2-}$
MW: 2371,76

Protected diphosphoinositol pentakisphosphate **13** (34.0 mg, 12.8 µmol, 1.0 eq.) was dissolved in DCM (2.0 mL), TFA (9.90 µL, 14.7 mg, 12.8 µmol, 10 eq.) was added and the reaction mixture was stirred for 15 min. The solvent was removed under high vacuum. The crude product was dissolved in MeCN and directly purified by an automated MPLC (Interchim-system C18aq-column, $H_2O$/MeCN with TEAA-buffer, 5mM) to avoid decomposition. The title compound **13a** (11.0 mg, 38.5 µmol, 30%) was isolated as TEAA-salt and colorless solid.

**$^1$H-NMR** (400 MHz, $D_2O$): δ = 7.42 − 6.83 (m, 44H), 5.45 − 4.89 (m, 28H), 2.29 − 2.24 (m, 33H) ppm. **$^{31}$P{$^1$H}-NMR** (162 MHz, $D_2O$): δ = -1.49 − -2.17 (m, 5P), -10.44 (d, *J* = 14.6 Hz), -11.85 (d, *J* = 14.4 Hz) ppm.

**Synthesis of $^{18}$O-labeled 4-PP-InsP$_7$ (14)**

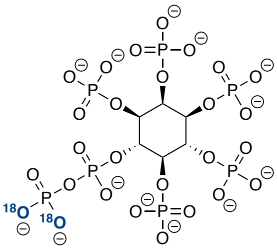

**14**

$C_6H_6O_{25}{}^{18}O_2P_7{}^{13-}$
MW: 730,91

${}^{18}O$-labeled diphosphoinositol pentakisphosphate **13a** (11.0 mg, 4.2 µmol, 1.0 eq.) was dissolved in DMF (1.0 mL) and piperidine (0.5 mL) was added. The resulting mixture was stirred for 1.5 h at rt. The mixture was diluted with $H_2O$ (20 ml) and directly purified by an automated SAX purification system (Äkta pure, HiTrap Capto™ Q ImpRes column, elution by an increasing gradient of $NH_4HCO_3$ buffer, 1M, pH 8.0). Fractions were analyzed by CE-MS measurement and the product containing fractions were combined and lyophilized. The title compound **14** (3.0 mg, 3.3 mmol, 78%) was obtained as a colorless solid.

**${}^{1}$H-NMR** (400 MHz, $D_2O$): δ = 5.05 − 4.97 (m, 1H), 4.71 − 4.63 (m, 1H), 4.63 − 4.53 (m, 1H), 4.40 − 4.25 (m, 1H) ppm. **${}^{31}$P{${}^{1}$H}-NMR** (162 MHz, $D_2O$): δ = 0.48 (s, 1P), -0.06 (s, 1P), -0.27 (s, 1P), -0.48 (s, 1P), -1.00 (s, 1P), -10.90 (d, *J* = 17.3 Hz), -11.85 (d, *J* = 18.2 Hz) ppm. **HRMS** (ESI) m/z for $C_6H_{17}O_{25}{}^{18}O_2P_7$ $[M]^{2-}$ : calcd. 370.9108 found 370.9115.

Compound 12: $^{1}$H-NMR (400 MHz, $CDCl_3$):

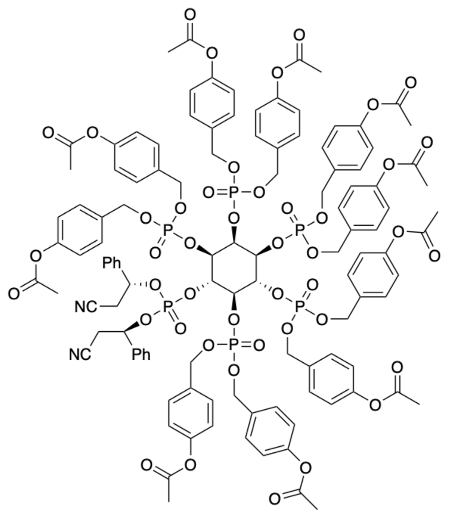


**12**

$C_{114}H_{112}N_2O_{44}P_6$
MW: 2399,96

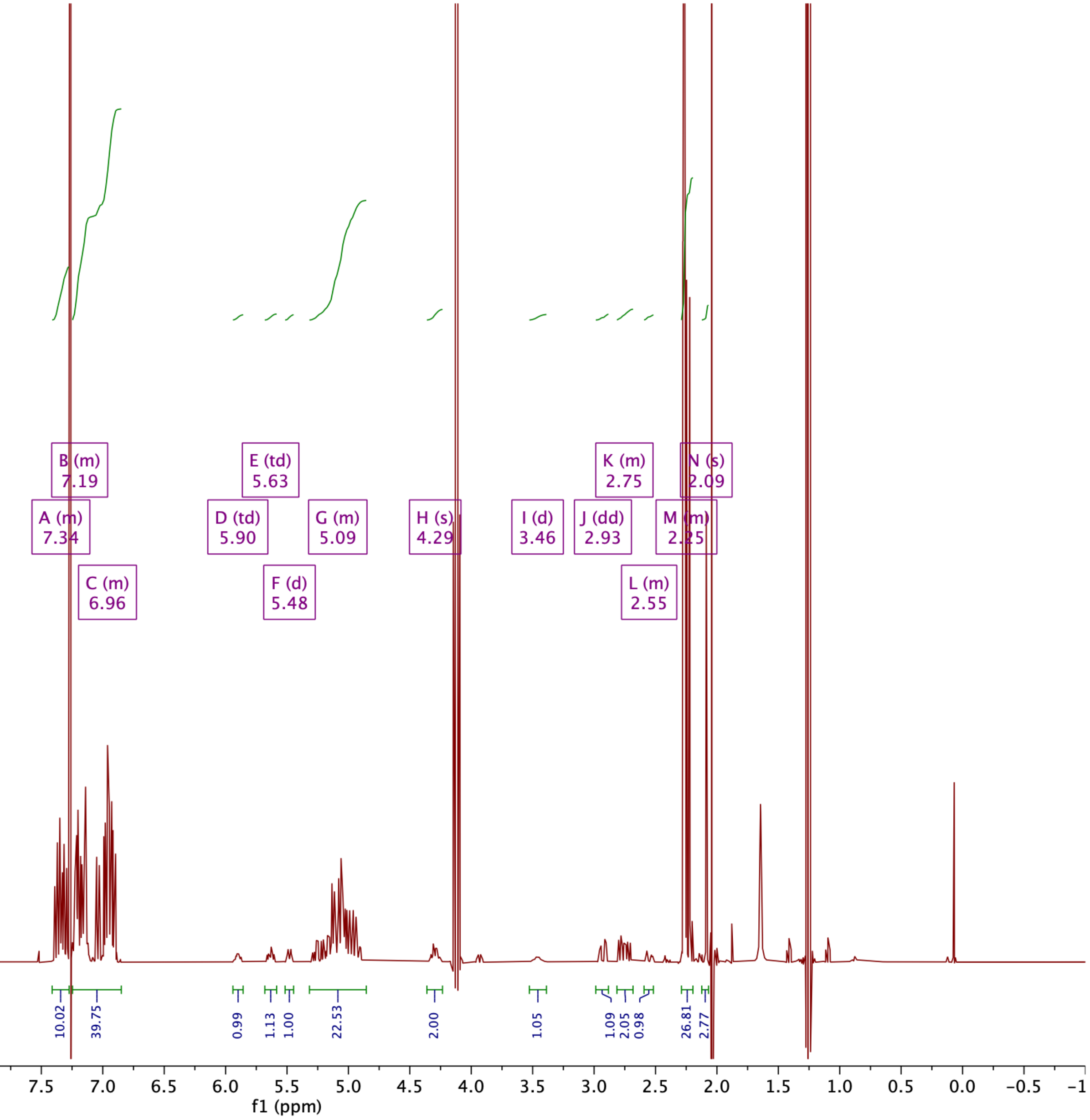

Compound 12: $^{31}$P-{$^{1}$H}NMR (162 MHz, $CDCl_3$):

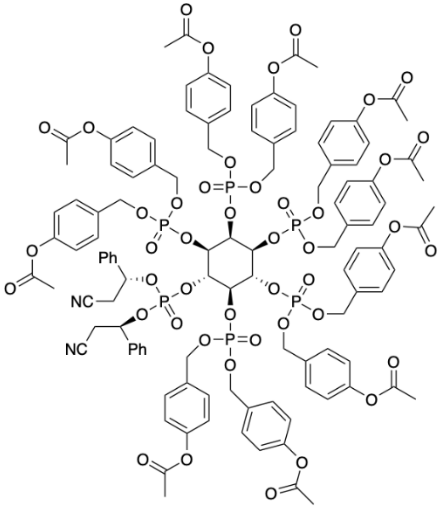

**12**

$C_{114}H_{112}N_2O_{44}P_6$
MW: 2399,96

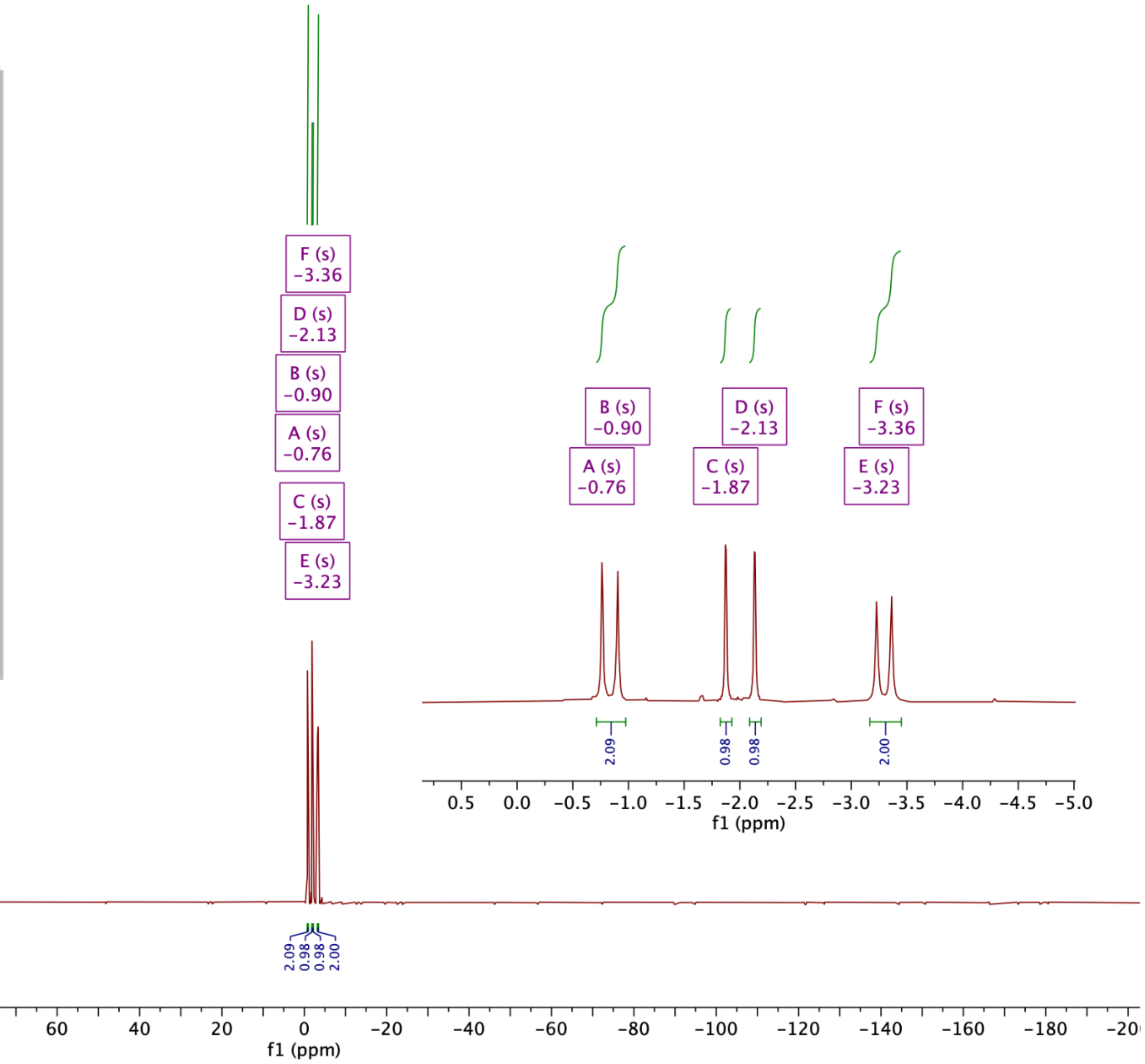

Compound 13a: $^{1}$H-NMR (400 MHz, $CDCl_3$):

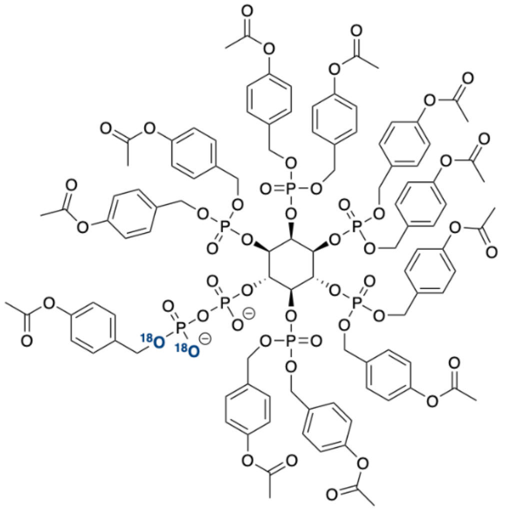

13a

$C_{105}H_{105}O_{47}{}^{18}O_2P_7{}^{2-}$
MW: 2371,76

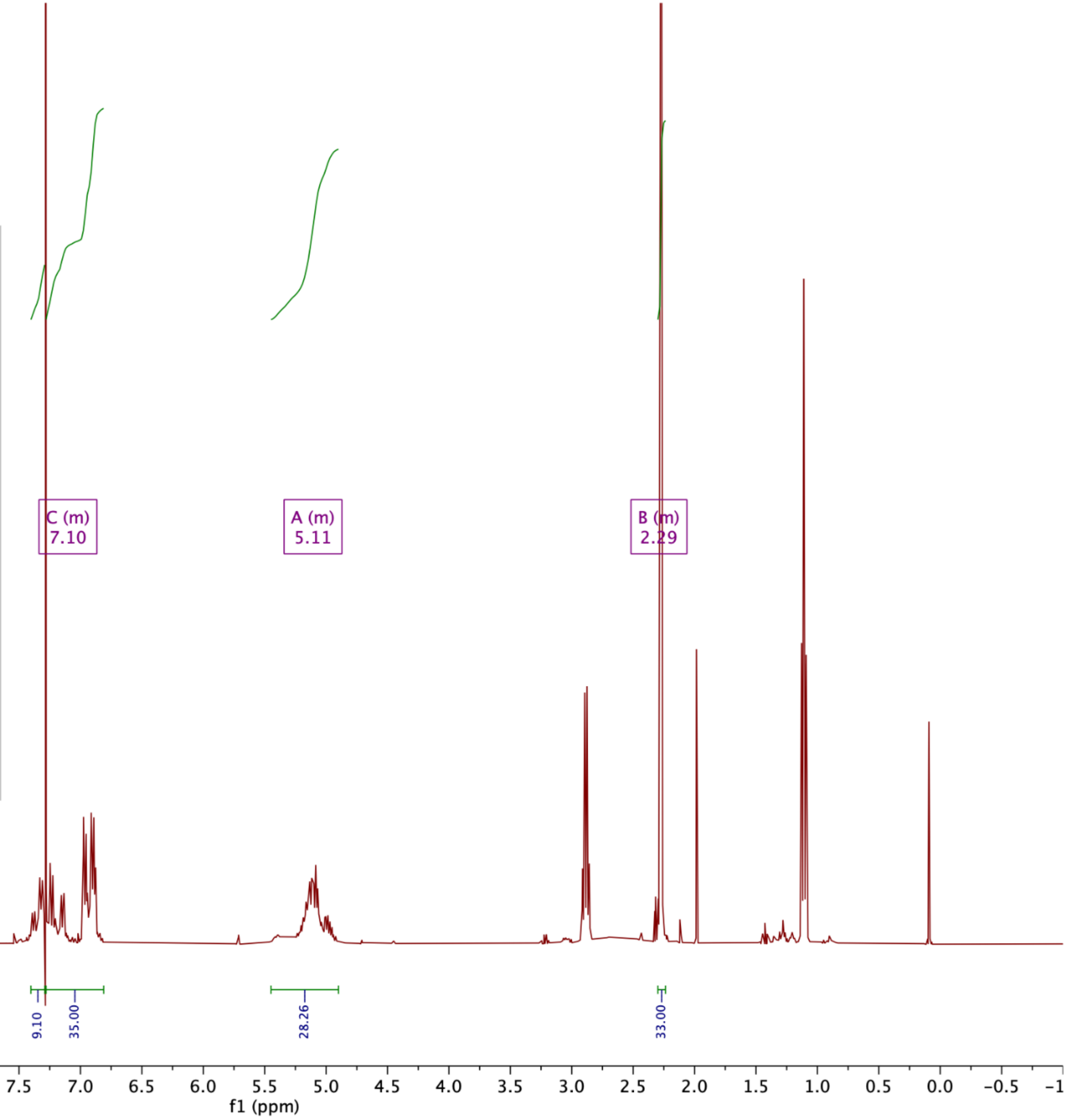

Compound 13a: $^{31}$P-{$^{1}$H}NMR (162 MHz, $CDCl_3$):

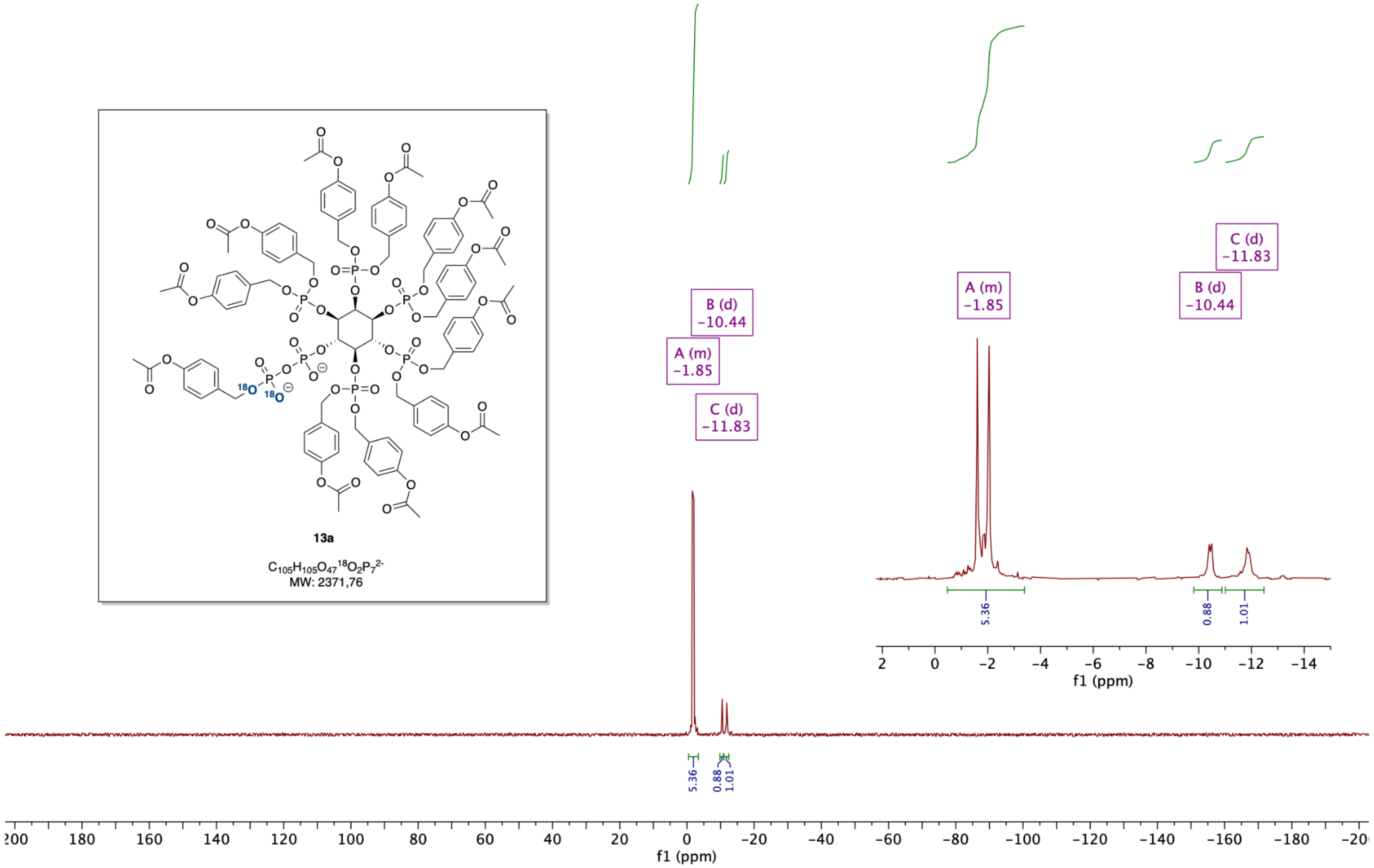

Compound 14: $^{1}H$-NMR (400 MHz, $D_2O$, contains phosphonoacetic acid as internal standard):

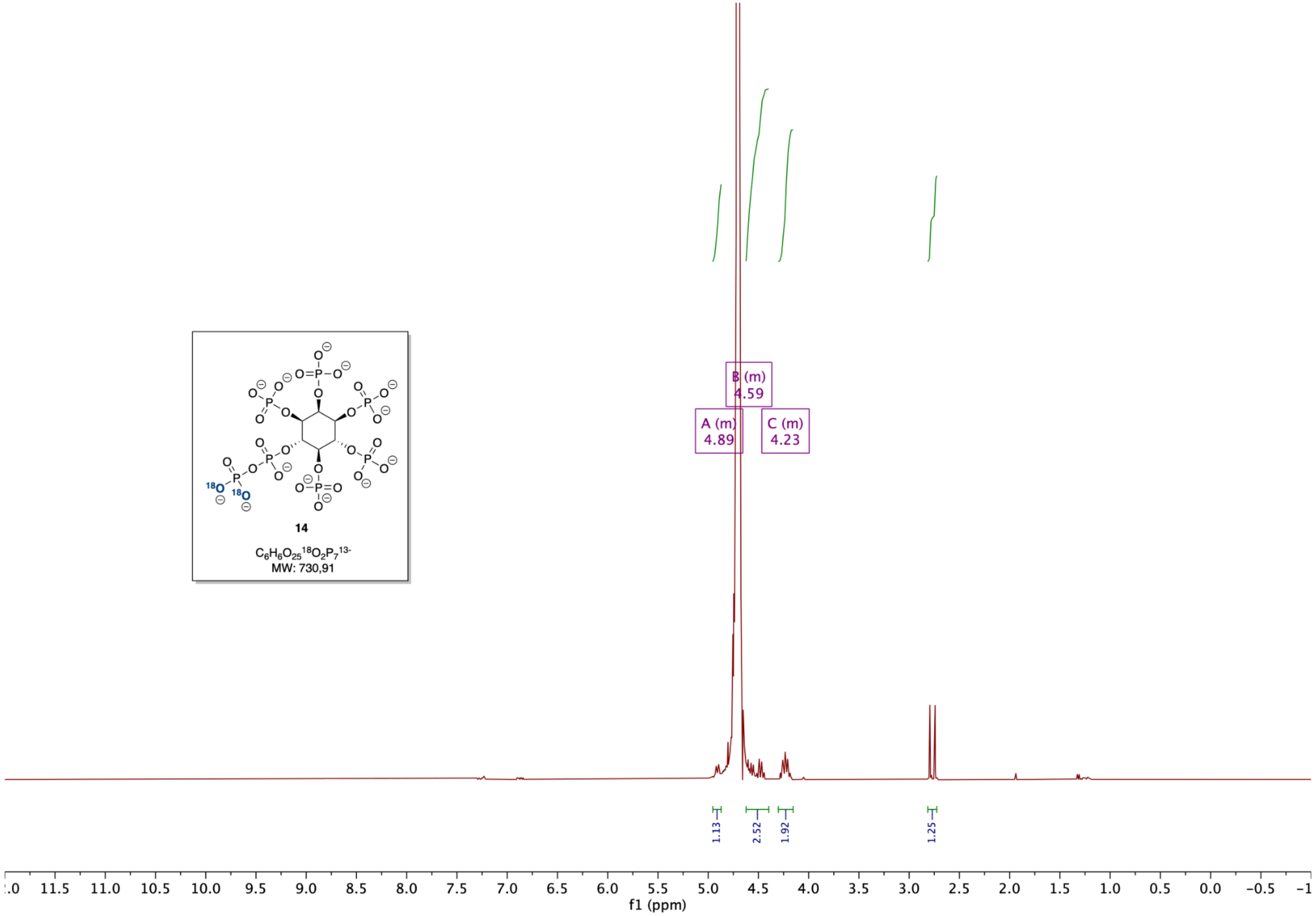

Compound 14: $^{31}P$-{$^{1}H$}NMR (162 MHz, $D_2O$, contains phosphonoacetic acid as internal standard):

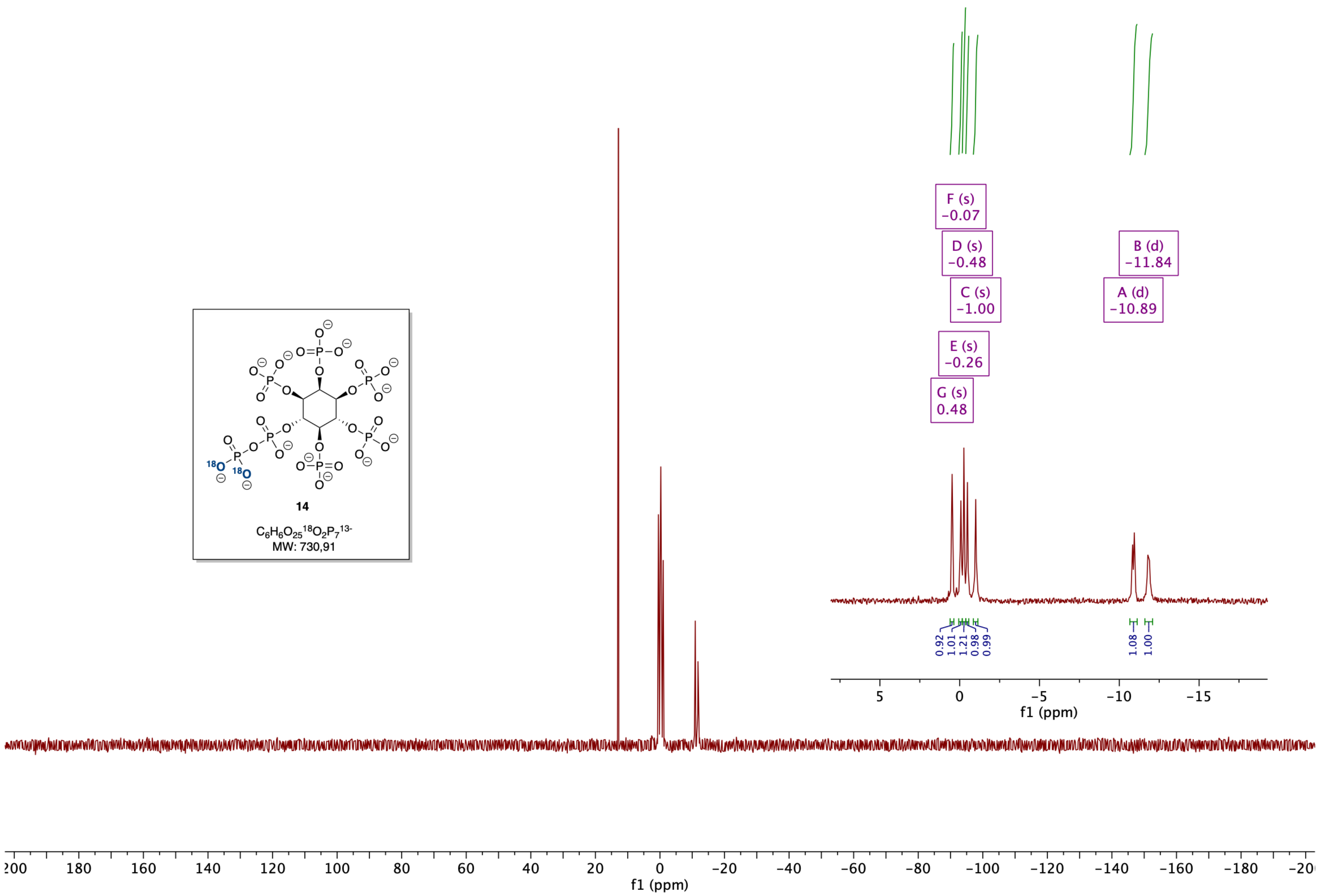

**Supporting References**

[1] Jessen, H. J.; Schulz, T.; Balzarini, J.; Meier, C. *Angew. Chemie Int. Ed.* **2008**, *47* (45), 8719–8722.
[2] Capolicchio, S.; Thakor, D. T.; Linden, A.; Jessen, H. J. *Angew. Chemie - Int. Ed.* **2013**, *52* (27), 6912–6916.
[3] T. M. Haas, S. Mundinger, D. Qiu, N. Jork, K. Ritter, T. Dürr-Mayer, A. Ripp, A. Saiardi, G. Schaaf, H. J. Jessen, *Angew. Chem. Int. Ed.* **2022**, *61*, e202112457.
